# Decoding Hierarchical Cell–Cell Communication in Spatial Multi-Omics with CellSTIC

**DOI:** 10.64898/2026.05.27.728114

**Authors:** Shuai Wang, Jiayin Wang, Jiayi Wang, Zhiyuan Yuan, Yungang Xu

## Abstract

Cell–cell communication coordinates tissue development, homeostasis, and immunity, yet defining signaling interactions within intact tissues remains challenging. Single-cell transcriptomics enables systematic ligand– receptor inference, but tissue dissociation removes spatial context and obscures local and region-specific signaling. Spatial transcriptomics and spatial multi-omics can recover communication in situ, although existing methods often incompletely integrate heterogeneous data or produce poorly interpretable ligand–receptor lists. Here we present CellSTIC, a framework that resolves cell–cell communication in spatial multi-omics as structured programs grounded in tissue architecture. CellSTIC integrates multimodal evidence from local neighborhoods and organizes interactions into a hierarchical semantic representation that remains traceable to underlying molecules. This enables analysis from individual ligand–receptor pairs to functional modules comparable across tissues, regions, and states. In simulations and diverse tissue datasets, CellSTIC recovered spatially coherent communication structures and domains, revealing immune, brain, developmental, and regenerative programs, and providing a general approach for generating mechanistic hypotheses in situ.

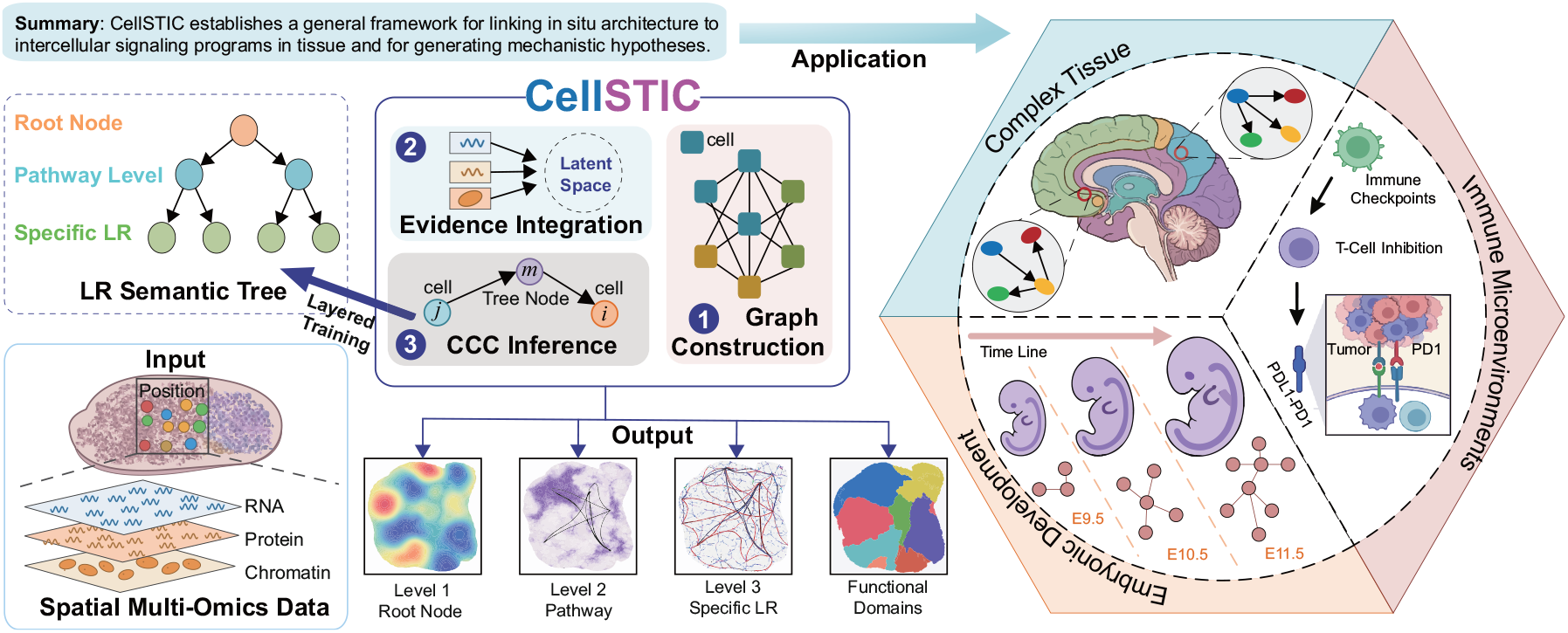

## 1 Introduction

Cell–cell communication (CCC) is a fundamental feature of multicellular life, enabling distinct cell populations to coordinate differentiation, tissue organization, and functional homeostasis through the exchange of molecular signals [1]. Among the many forms of intercellular communication, ligand–receptor (LR) interactions are one of the most prevalent and experimentally tractable mechanisms, because they provide a direct link between signal production, signal reception, and downstream cellular responses. Large-scale single-cell atlases have shown that most cells simultaneously express numerous ligands and receptors, giving rise to densely interconnected signaling networks that shape tissue organization and function [2].

When such LR-mediated signaling becomes imbalanced, rewired or aberrantly interpreted within the tissue microenvironment, intercellular communication networks are correspondingly remodeled, contributing to processes such as tumor progression, immune evasion, and therapeutic resistance [3]. Identifying, in situ, which cell populations communicate, and through which key signaling axes, is therefore central to understanding both tissue homeostasis and disease-associated reprogramming, and is also relevant to the discovery of actionable therapeutic targets [4, 5]. Before spatially resolved profiling became widely available, dissociation-based single-cell RNA sequencing (scRNA-seq) and its associated computational methods provided the main route for CCC inference. These approaches substantially improved the resolution of communication analysis, from cell-type-level co-expression to finer-grained interaction modeling. For example, Scriabin extended inference to the level of individual cell pairs, whereas methods such as DcjComm sought to connect LR interactions with downstream regulatory evidence [6, 7]. However, because dissociation disrupts tissue architecture and local neighborhood relationships, these approaches cannot reliably distinguish bona fide proximal communication from signals driven by covariance in cell composition or cell state, limiting the identification of spatially restricted communication programs and the generation of testable mechanistic hypotheses.

Recent advances in spatial transcriptomics [8, 9] and spatial multi-omics technologies [10, 11] provide an opportunity to overcome this limitation. By preserving spatial coordinates while measuring multiple molecular layers in the same tissue section, these platforms make it possible to study communication together with tissue architecture in situ and to localize signaling events to defined microenvironments and anatomical units [12, 13]. Considerable progress has already been made in methods for CCC inference from spatial data. COMMOT, for example, introduces spatial constraints and competition among ligands and receptors within an optimal transport framework [14]. CellChat and related methods organize intercellular signaling into pathway-level communication networks and support comparisons across biological contexts [15, 16]. Deep-learning-based methods such as CellNEST further explore more complex spatial communication structures, including relay-like interactions [17]. Together, these advances have established spatially informed CCC analysis as an important direction for dissecting tissue organization and function.

Despite this progress, two major challenges remain, particularly for spatial multi-omics data. First, the presence of multiple modalities does not automatically translate into stronger or more reliable communication evidence. Differences in information content, noise structure, measurement bias, and resolution across modalities mean that naïve fusion can introduce scale mismatch and bias accumulation, resulting in unstable inference in which strong modalities dominate and weaker but informative modalities are diluted [18, 19]. Second, most existing methods still represent CCC primarily as flattened lists of LR pairs. Although useful as a starting point, such outputs often produce large numbers of candidate interactions that are difficult to compare across regions, conditions, or developmental stages, and they do little to organize dispersed signals into higher-order functional programs [20]. As a result, an important conceptual gap remains between identifying individual interaction candidates and deriving coherent, interpretable mechanistic narratives from spatial multi-omic data.

We therefore reasoned that CCC analysis in spatial multi-omics should address both evidence integration and knowledge representation. At the evidence level, spatial proximity and cross-modal complementarity should be incorporated explicitly so that inferred communication is supported more consistently by multiple sources of observation. At the representation level, individual LR interactions should be organized into structured, biologically interpretable units that allow communication patterns to be compared, traced, and summarized across tissues and contexts. Hierarchical organization provides a natural framework for this goal, because it can group related interactions into modules while preserving links between higher-order programs and the underlying molecular evidence. Here, we present CellSTIC (Spatial and Tree-Informed Communication), a framework that views cell–cell communication in spatial multi-omics as a feature of tissue organization rather than isolated ligand–receptor events. By integrating multimodal signals from local neighborhoods and resolving them within a hierarchical representation, CellSTIC connects molecular interactions to higher-order communication programs while preserving their spatial context. This design creates a scalable and interpretable foundation for studying how signaling is organized across tissues, regions, and developmental and regenerative states. More broadly, CellSTIC offers a framework for linking tissue architecture to intercellular communication and for generating mechanistic insight from spatial multi-omic data.

## 2 Results

### 2.1 CellSTIC Framework

CellSTIC was developed to infer spatially resolved cell–cell communication within a structured biological context, rather than treating ligand–receptor pairs as isolated prediction targets. In spatial tissues, communication signals are organized across nested functional levels and are closely linked to spatial domains. A flat interaction label space, therefore, limits biological interpretation and obscures higher-order communication programs. CellSTIC addresses this challenge as an end-to-end spatial multi-omics framework that jointly performs communication inference and spatial functional-domain identification within a tree-structured semantic system (Fig. 1**a**). Starting from spatial multi-omics profiles and coordinates, CellSTIC operates directly on capture units, referred to here as spots, and accommodates diverse combinations of molecular modalities. The framework is designed to infer spatially resolved communication strengths and interaction identities across multiple semantic resolutions while simultaneously deriving stable tissue domains.

**Fig. 1.**
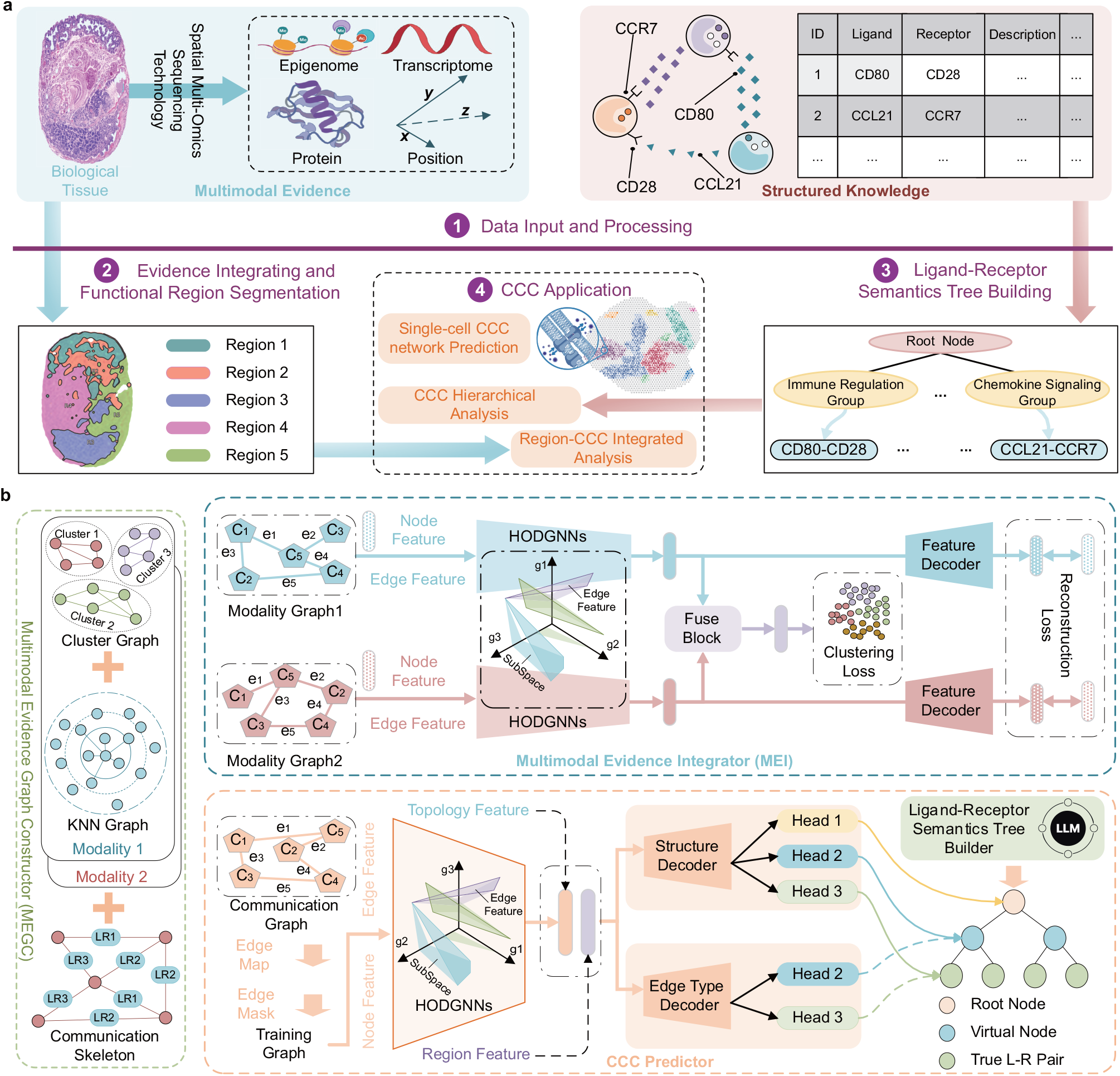
Overview of the CellSTIC framework. **a**, Workflow of CellSTIC. (1) Spatial multi-omics data and structured ligand–receptor knowledge are used as input. (2) Multimodal molecular evidence and spatial information are integrated for functional region segmentation. (3) A ligand–receptor semantics tree is constructed. (4) Cell–cell communication is then analyzed across hierarchical semantic levels and spatial regions. **b**, Architecture of the main modules and training strategy. The Multimodal Evidence Graph Constructor (MEGC) builds modality-specific graphs and a ligand–receptor-mediated spatial backbone graph. The Multimodal Evidence Integrator (MEI) learns unified representations from multimodal inputs through modality-specific encoding and fusion, together with clustering and reconstruction objectives. The CCC predictor models communication on the spatial backbone graph with edge-semantic-aware graph learning and is trained using an edge masking and reconstruction strategy under the guidance of the ligand–receptor semantics tree. The model outputs, for each spatially proximal cell pair, communication strength and interaction-type predictions across multiple semantic levels. Both MEI and the CCC predictor use the Hyperplane Orthogonal Decomposition Graph Neural Network (HODGNN) as the encoding backbone.

After preprocessing, CellSTIC formulates the task as graph learning under spatial constraints (Fig. 1**b**). It first constructs a spatial communication graph using tissue geometry and ligand–receptor evidence, thereby restricting candidate interactions to spatially plausible events. This graph is then refined with multimodal information from local neighborhoods and broader tissue structure, allowing communication inference to retain fine-grained heterogeneity while remaining consistent with tissue-scale organization. CellSTIC further implements this step through HODGNN, an edge-semantic-aware message-passing operator that incorporates spatial distance, directionality, and interaction strength into graph propagation. This enables joint updates of node and edge representations while preserving spatial cues during representation learning. To support multimodal integration, CellSTIC learns a unified latent representation that captures cross-modal consensus together with modality-specific variation. A second-stage communication-specific fine-tuning step then reshapes this latent space towards interaction-relevant signals while preserving the multimodal tissue structure learned in the first stage. The resulting embeddings provide a shared representation for communication inference and domain identification, enabling the two tasks to reinforce one another.

A central feature of CellSTIC is its hierarchical representation of interaction types. Instead of modelling ligand–receptor pairs as independent labels, CellSTIC organizes interactions into a traceable semantic tree and performs coarse-to-fine prediction through self-supervised training (Fig. 1**b**). This design enables multi-resolution communication outputs, from broad functional classes to specific ligand–receptor identities, and improves inference for weak or sparse signals by using higher-level semantic structure to constrain fine-grained predictions. To enhance stability and interpretability, the hierarchy is further supported by class balancing, semantic organization, and prior biological knowledge (Extended Data Fig. 1**b**). Together, these components allow CellSTIC to couple spatial multimodal evidence with hierarchical interaction semantics, representing communication as structured, spatially resolved biological programs rather than fragmented lists of candidate interactions. Ablation, sensitivity, and scalability analyses are provided in Supplementary Fig. S1 and Supplementary Notes S5 and S6.

### 2.2 CellSTIC Outperforms Existing Methods in Cell–Cell Communication Prediction and Spatial Region Identification

We evaluated CellSTIC on a unified multimodal CCC benchmark and compared it with COMMOT [14], CellNEST [17], Scriabin [6], and DcjComm [7]. scMultiSim [21] simulations provided spatial coordinates, cell-type composition, RNA–ATAC profiles, and ground-truth communication links. All methods were assessed on matched replicates, inputs, and preprocessing settings using recommended or default parameters. Results were aggregated across eight replicates, with Figs. 2**a**–**d** showing representative visualizations from re1.

**Fig. 2.**
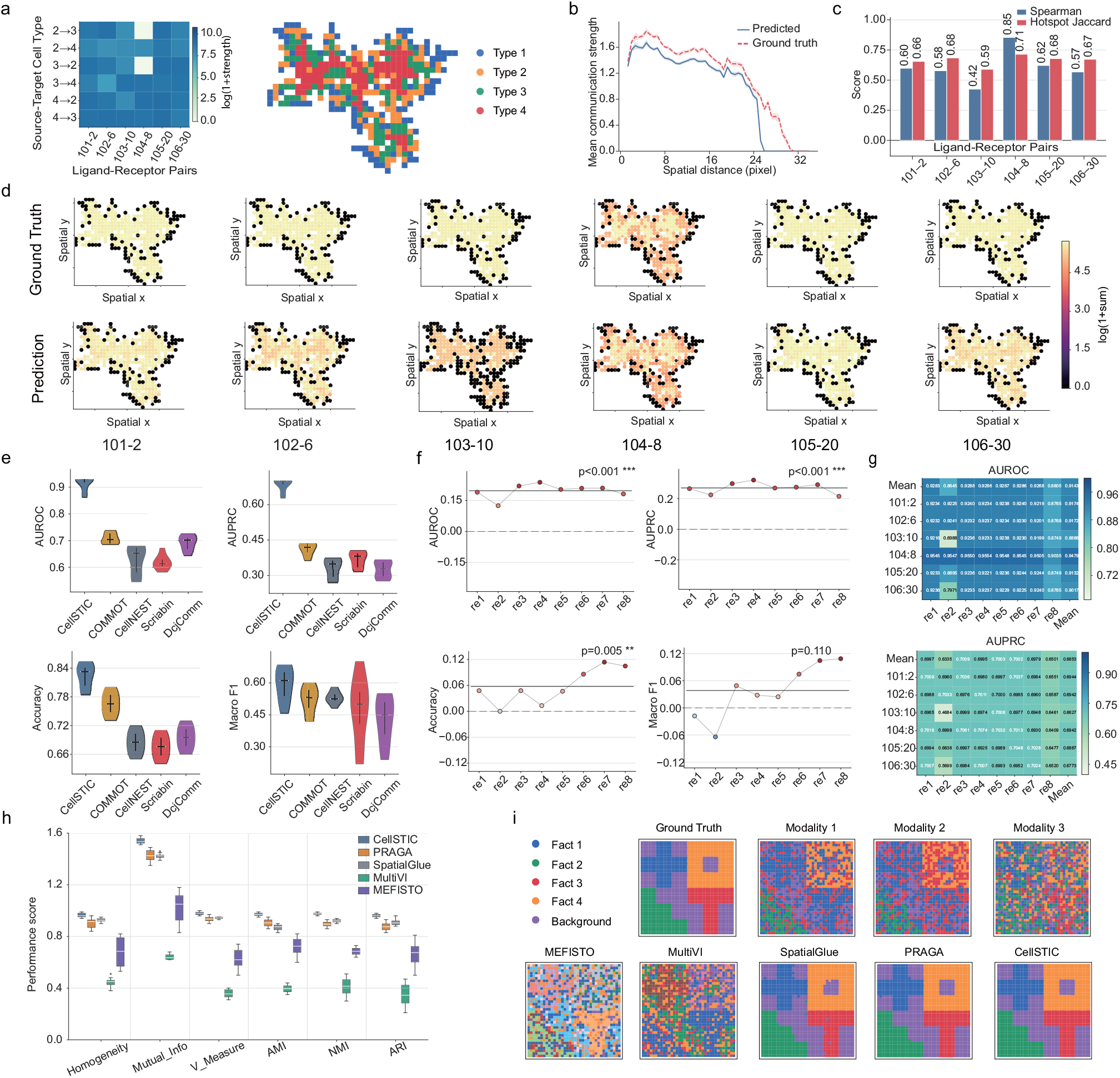
Benchmarking on simulated multimodal spatial data shows accurate cell–cell communication prediction and robust spatial region identification. **a**, Predefined ligand–receptor-specific communication strengths, cell-type labels, and spatial partitions enable joint evaluation of CCC inference and region identification. Left, aggregated region-to-region strengths shown as log(1+sum); right, ground-truth spatial partition. **b**, Mean predicted communication strength as a function of spatial distance, compared with the ground truth, showing recovery of distance-dependent decay. **c**, Recovery of spot-level communication patterns for representative ligand–receptor pairs, quantified by the Spearman correlation of spot scores and the Jaccard overlap of hotspot regions defined by the top 10% highest-scoring spots. **d**, Representative ground-truth and predicted spatial communication maps for individual ligand– receptor pairs, showing accurate reconstruction of pair-specific signaling patterns. **e**, Performance of CellSTIC and baseline methods for binary CCC edge prediction, evaluated by AUROC, AUPRC, Accuracy, and Macro-F1. **f**, Improvement of CellSTIC over the strongest baseline across simulation replicates for the four CCC prediction metrics (Δ = ours *™*best baseline). **g**, Reproducibility of CCC prediction across replicates, shown by pairwise performance matrices for AUROC and AUPRC. **h**, Performance of CellSTIC and baseline methods for spatial region identification, evaluated by Homogeneity, Mutual Information, V-Measure, Adjusted Mutual Information (AMI), Normalized Mutual Information (NMI), and Adjusted Rand Index (ARI). **i**, Representative spatial partitions from the simulated benchmark, comparing the ground truth, unimodal observations, and multimodal baseline methods with CellSTIC.

In re1, CellSTIC accurately reconstructed the non-uniform communication landscape across cell-pair and ligand–receptor (LR) dimensions, consistent with the underlying spatial organization of four cell types (Fig. 2**a**). Predicted communication strengths showed clear distance-dependent decay and closely matched the ground truth across spatial scales (Fig. 2**b**). At the spot level, CellSTIC achieved strong rank concordance with ground truth (Spearman correlation, 0.42–0.85) and accurately localized communication hotspots, defined as the top 10% high-scoring spots (Jaccard index, 0.58–0.68; Fig. 2**c**). Spatial maps further confirmed recovery of both hotspot positions and LR-specific intensity gradients (Fig. 2**d**), indicating that CellSTIC captures tissue-structured communication rather than spatially unrelated noise. At the cell-pair edge level, CellSTIC consistently outperformed baseline methods across re1–re8 (Fig. 2**e**), achieving AUROC 0.92 ±0.03, AUPRC 0.68 ±0.03, accuracy 0.82 ±0.03, and macro-F1 0.56 ±0.12. Relative to the strongest baseline in each replicate, CellSTIC showed consistent gains in AUROC, AUPRC, and accuracy (Δ ≈0.1–0.3; Fig. 2**f**). Performance remained broadly stable across the replicate × LR-pair space, with reduced variability for most LR pairs (Fig. 2**g**). Together, these results show that CellSTIC recovers spatially constrained communication patterns, improves edge-level prediction, and provides robust performance across replicates and LR pairs.

We next evaluated whether CellSTIC could recover tissue-structure-consistent spatial domains, providing a robust context for downstream communication analysis. Using the Spatial Multimodal Dataset following the SpatialGlue evaluation setting [18], spatial structure was generated with ggblocks and non-negative spatial factorization as described by Townes et al. [22]. The dataset includes three simulated modalities, namely RNA, ADT, and ATAC, together with ground-truth spatial domains for supervised assessment. Compared with representative multimodal integration methods, including MEFISTO [23], MultiVI [24], PRAGA [25] and SpatialGlue, CellSTIC achieved consistently higher scores with lower variability across homogeneity, MI, V-measure, AMI, NMI and ARI (Fig. 2**h**). CellSTIC also produced clearer spatial domain boundaries and reduced cross-domain mixing in visualizations (Fig. 2**i**). Similar trends were observed in real human skin and tonsil samples, with full results provided in Supplementary Note S7 and Supplementary Fig. S2. Together, these benchmarks indicate that CellSTIC not only improves communication prediction but also preserves spatially coherent interaction patterns and tissue-consistent domain organization.

### 2.3 CellSTIC Resolves Hierarchical Communication Programs Defining Immune Microenvironments in the Human Lymph Node

To test whether CellSTIC resolves hierarchical communication programs in complex immune tissue, we applied it to a human lymph node spatial multi-omics dataset. The lymph node contains highly organized stromal niches, chemokine gradients, and regionally positioned immune populations that together define local microenvironments. Although RNA and ADT each revealed spatial heterogeneity, their partitioning patterns were only partly concordant (Extended Data Fig. 2**b**). By integrating both modalities, CellSTIC recovered spatial regions that were more continuous and more consistent with known lymph node architecture (Fig. 3**a**). These regions reflected distinct cellular ecological contexts, with marked shifts in B lineage, Helper and Regulatory T, Cytotoxic and innate-like T, Myeloid, NK and ILC, Plasma lineage, Erythroid and MK, and Stromal and non-immune compartments (Fig. 3**b**). Thus, CellSTIC provided an anatomically grounded basis for hierarchical communication analysis.

**Fig. 3.**
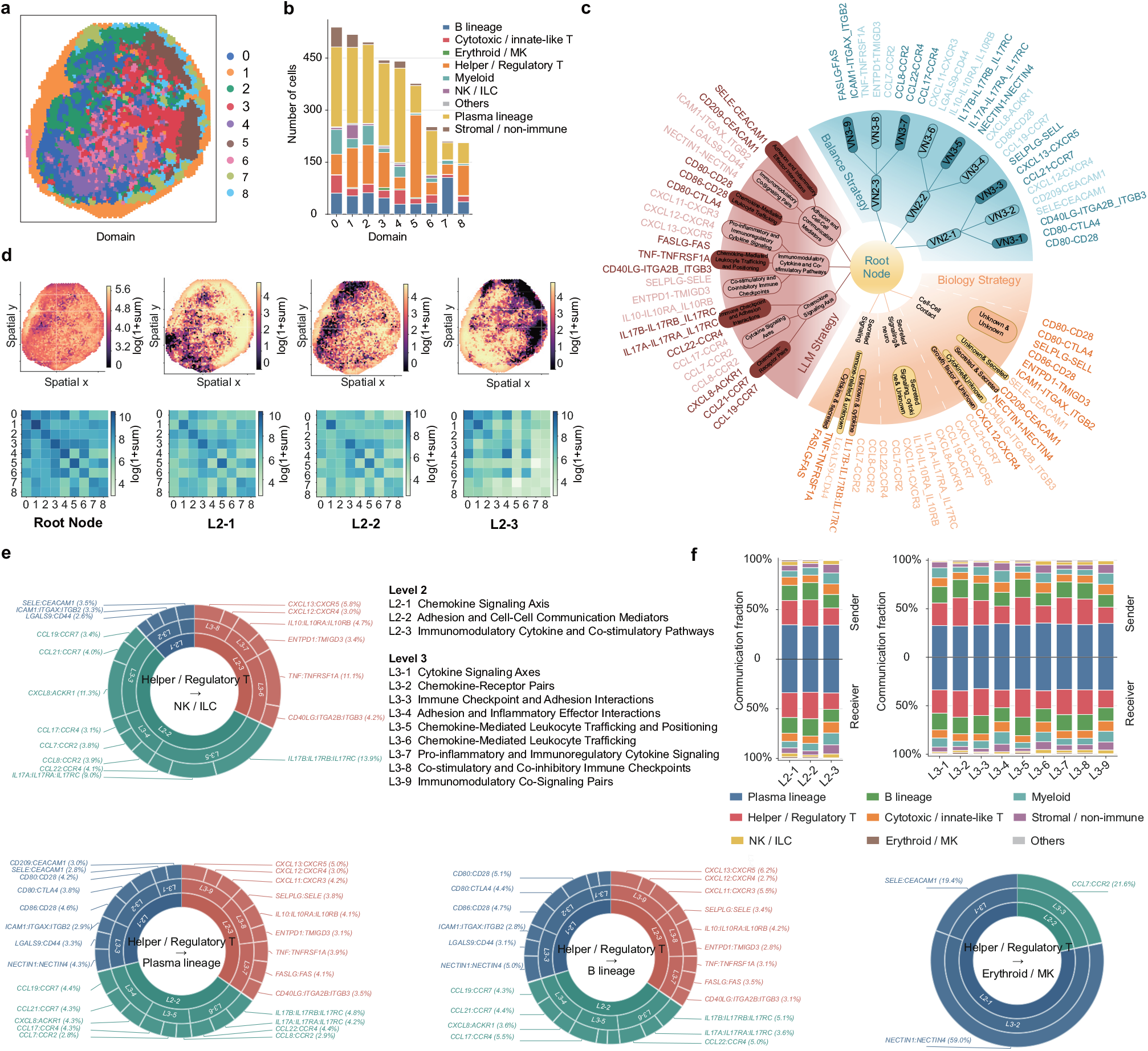
LLM-guided ligand–receptor semantic hierarchy reveals multiscale spatial communication in the human lymph node. **a**, CellSTIC identifies spatially coherent communication domains in the human lymph node. **b**, The inferred communication domains correspond to distinct immune microenvironments with characteristic cell-type compositions. **c**, CellSTIC learns a multiscale ligand–receptor semantic hierarchy under balanced constraints, biological priors, and LLM-guided semantic supervision, enabling interpretable organization of communication programs without cell-type labels. **d**, CCC exhibits spatially structured heterogeneity across the tissue and across hierarchy levels, with the root node capturing global communication structure and second-level nodes resolving region-specific programs. Per-cell communication intensity was projected onto spatial coordinates after log(1 + *x*) transformation. **e**, Post hoc summarization of learned hierarchical nodes shows that communication from Helper/Regulatory T cells to Erythroid/MK, NK/ILC, B-lineage, and Plasma-lineage compartments is selectively enriched in distinct branches of the hierarchy. For each cell-type pair, node-level communication strength was defined as the total probability mass of ligand–receptor pairs assigned to each node. **f**, Sender and receiver cell-type composition across hierarchical communication nodes at the second and third levels.

We next organized 26 candidate ligand–receptor pairs related to lymph node homeostasis and immune regulation into coarse-to-fine semantic hierarchies (Fig. 3**c**). Individual ligand–receptor pairs formed the leaf nodes, whereas internal nodes aggregated related interactions into broader functional modules. Across hierarchy construction strategies, the resulting trees transformed a flat interaction list into an interpretable modular structure, with downstream analyses focused primarily on the LLM-guided hierarchy. Canonical chemotactic signals, including CCL19-CCR7 and CCL21-CCR7, together with CXCL8-ACKR1, were grouped within a chemokine-related branch, linking individual interactions to a higher-order chemokine signalling axis. This organization confirmed that migration- and positioning-associated interactions can be recovered as coherent functional modules.

Projecting communication strengths from different hierarchy levels onto tissue space revealed multiscale spatial heterogeneity (Fig. 3**d**). Root-level signals were broadly distributed, consistent with a shared communication background shaped by stromal support, chemokine gradients, and compartmentalized immune organization. Lower hierarchy levels showed more localized and region-biased patterns, indicating that chemotaxis, adhesion, and immune regulatory programs are preferentially deployed in specific microenvironments. Region-to-region communication matrices showed the same trend, with root nodes capturing a shared interaction scaffold and lower-level modules decomposing this scaffold into differentiated regional interaction programs. Post hoc attribution further revealed that communication from Helper and Regulatory T cells to different immune compartments was partitioned into distinct branch-restricted programs (Fig. 3**e**). Interactions with B-lineage and Plasma-lineage compartments spanned broader regions of the semantic tree, including chemokine-related and immunomodulatory branches, consistent with composite programs involving positioning, helper activity, and immune regulation. Representative interactions included CXCL13-CXCR5, CCL19-CCR7, CCL21-CCR7, IL10-IL10RA_IL10RB, and TNF-TNFRSF1A, in line with the role of T cell-derived help in shaping B-cell fate and plasma-cell output [26, 27]. By contrast, communication with NK and ILC, and especially Erythroid and MK compartments, was confined to narrower hierarchy sectors dominated by chemotactic or adhesion-related modules [28, 29]. Thus, hierarchical semantic integration distinguished broadly distributed composite programs from more compartment-specific terminal effector programs.

Finally, sender-receiver communication composition across hierarchy levels revealed a stable tissue-level scaffold dominated by Plasma lineage and Helper and Regulatory T cells at the second level (Fig. 3**f**). At the third level, this scaffold was largely preserved, but individual submodules showed interpretable shifts: chemokine-related branches had greater contributions from B lineage and Helper and Regulatory T compartments, whereas cytokine, co-signalling and adhesion-related branches recruited broader contributions from Myeloid, Cytotoxic and innate-like T, and Stromal and non-immune compartments (Fig. 3**f**). At the leaf-node level, most ligand–receptor pairs retained the participation structure of their parent modules while preserving local molecular preferences (Extended Data Fig. 2**c**). Together, these results show that CellSTIC recovers known principles of lymph node organization and resolves how shared communication scaffolds diversify into compartment-specific effector programs across hierarchical semantic levels.

### 2.4 CellSTIC Dissects Spatially Embedded Communication Programs to Reveal Region-Specific Signaling in the Mouse Brain

To test whether CellSTIC resolves the spatial embedding of communication programs within fine tissue architecture, we applied it to a mouse brain spatial multi-omics dataset. The brain provides a stringent setting for spatial communication inference because it contains highly heterogeneous cell populations, fine-grained anatomical organization, and interactions constrained by local microenvironments. The original tissue image, cell-type composition, and modality-specific inputs are shown in Extended Data Figs. 3**a, b**. By integrating multimodal information within a unified spatial coordinate system, CellSTIC identified 15 spatial domains (Fig. 4**a**). These domains were broadly consistent with known neuroanatomical organization, covering cortical laminar territories, striatum-related regions, ventricle-adjacent areas, and white-matter tracts. This partitioning preserved large-scale tissue continuity while resolving local transition zones and finer structural features, providing an anatomically constrained framework for downstream communication analysis.

**Fig. 4.**
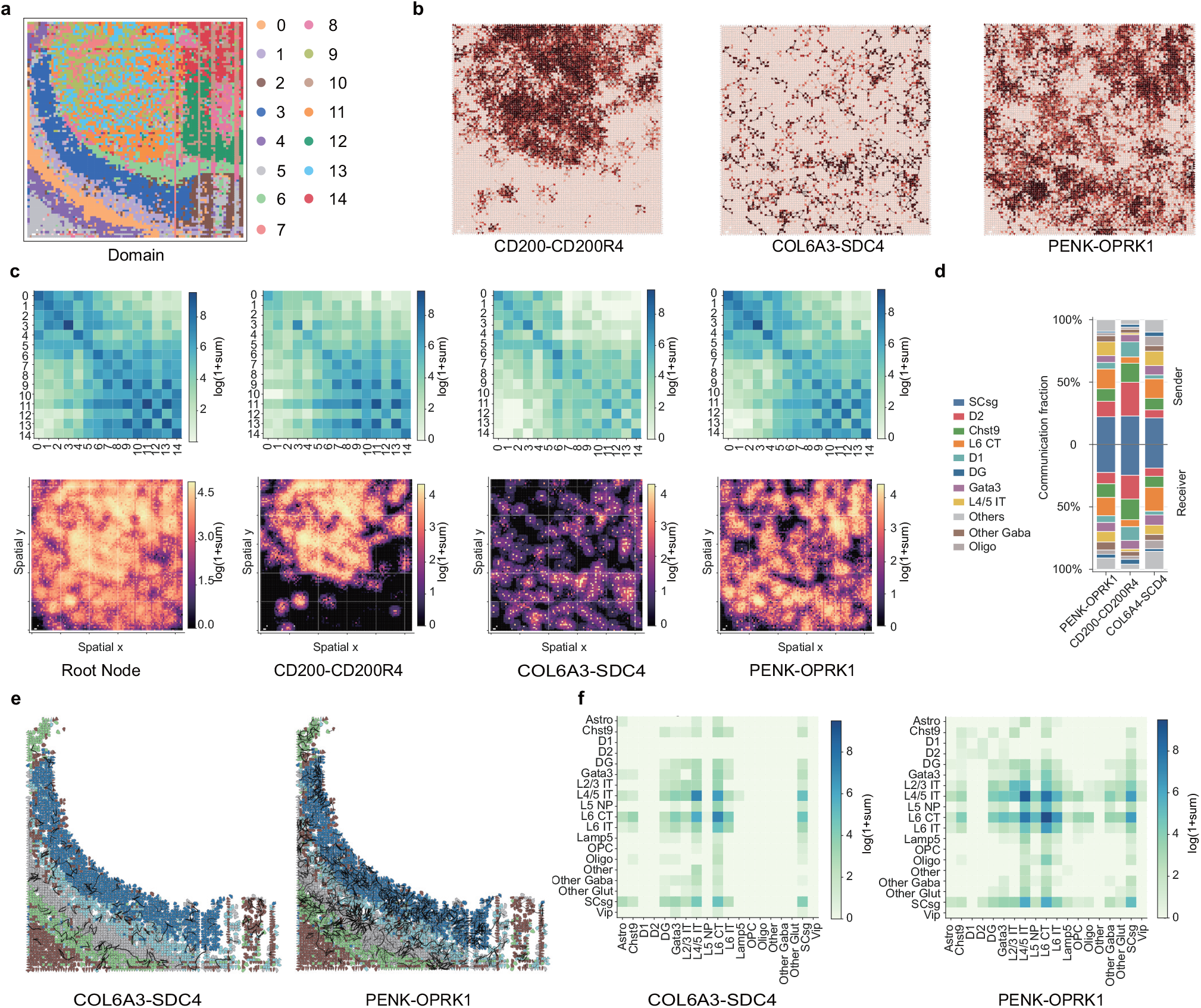
High-resolution spatial domains reveal region-specific cell–cell communication axes in the mouse brain. **a**, Spatial domains inferred from integrated spatial multi-omics analysis, identifying 15 domains across the tissue section. **b**, Spatial maps of inferred cell–cell communication activity for the three representative ligand–receptor pairs analysed in this study: CD200-CD200R4, COL6A3-SDC4, and PENK-OPRK1. **c**, Comparison of communication strength at the spatial-domain level. Top, spatial distributions of global CCC activity and of the three representative ligand–receptor pairs. Bottom, corresponding domain-by-domain communication heat maps. **d**, Major sender and receiver cell types contributing to each ligand–receptor pair, shown as communication fractions. **e**, Hierarchical visualization of CCC networks across spatial-domain boundaries. **f**, Cell-type-resolved sender–receiver communication heat maps for the corresponding boundary regions in **e**.

Within this spatial framework, we examined three representative communication axes, CD200-CD200R4, COL6A3-SDC4, and PENK-OPRK1. All three showed clear regional preference rather than uniform activity across the section (Fig. 4**b**). CD200-CD200R4 signals were concentrated in upper and central striatum-related regions, consistent with experimental evidence for spatially heterogeneous CD200-CD200R signalling across brain compartments [30]. COL6A3-SDC4 showed weaker activity and was enriched mainly at transitions between striatum-related and ventricle- or white-matter-adjacent regions. PENK-OPRK1 displayed broader multifocal hotspots, consistent with recent imaging studies showing subregion-specific endogenous opioid signalling dynamics in the brain [31]. These patterns were also reflected at the inter-domain level. CD200-CD200R4 interactions were largely confined to a limited subset of striatum-related domains, COL6A3-SDC4 showed weaker and more diffuse connectivity, and PENK-OPRK1 exhibited broader cross-domain interactions, suggesting a potential role in coupling distinct tissue compartments (Fig. 4**c**). Sender-receiver composition further differed across axes (Fig. 4**d**), indicating that partially overlapping communication programs coexist within the same tissue section and are shaped jointly by local proximity and tissue compartmentalization, in line with strong proximity effects reported in recent spatial brain atlases [32].

Global domain-level comparisons did not fully explain communication heterogeneity within individual regions. We therefore performed community detection on inferred CCC networks within domains 9 and 11 (Extended Data Fig. 4). Although both were assigned to coherent macroscopic spatial domains, CellSTIC further resolved each into spatially contiguous but molecularly distinct local communication communities. Domain 9 contained five subcommunities with 327, 174, 143, 103, and 80 spots, which were partially separated yet connected in UMAP space, indicating an internally structured organization rather than a uniform local niche (Extended Data Figs. 4**a, b**). Differential-feature analysis showed distinct molecular programs, including relative enrichment of Zcchc5, Myoc, and Pde1d in one group and increased Scarf1, Flt1, and Zeb2 in another (Extended Data Figs. 4**c, d**). Domain 11 was subdivided into three subcommunities with 429, 280, and 88 spots, showing clearer separation in both physical space and low-dimensional embedding, consistent with more discrete internal organization. We next extended the analysis to cross-region and boundary-associated communication. CCC networks for COL6A3-SDC4 and PENK-OPRK1 were preferentially organized along interfaces between laminar cortical territories and striatum-related regions, with additional extension toward ventricle- and white-matter-adjacent areas (Fig. 4**e**). COL6A3-SDC4 remained more confined to boundary and transition zones, whereas PENK-OPRK1 spanned broader areas, indicating distinct roles in cross-compartment coupling. Cell-type-resolved sender-to-receiver heat maps showed that both axes were concentrated in deep-layer excitatory neuronal populations, particularly L4/5 IT, L5 NP, L6 IT, and L6 CT-related combinations (Fig. 4**f**). Together, these results show that CellSTIC links tissue architecture, cell identity, and signalling activity within a common spatial framework, spanning domain identification, region-level communication mapping, intraregional microstructure dissection, and boundary-associated communication analysis.

### 2.5 CellSTIC Unveils Temporal Communication Programs Defined for Organ-Specific Development and Injury-Induced Regeneration

To test whether CellSTIC distinguishes distinct modes of temporal communication remodelling, we applied it to three complementary settings: mouse embryonic brain development, mouse embryonic liver development and axolotl telencephalon development versus injury-induced regeneration (Fig. 5**a**). The mouse series covered E9.5 to E16.5, capturing tissue regionalization, cell-state diversification and organ-level spatial constraints [33], whereas the axolotl datasets enabled comparison between normal telencephalon development and regeneration after injury [34]. Across these systems, cell-number dynamics already suggested distinct biological regimes, including rapid early expansion in the mouse brain, delayed but accelerated expansion in the liver, and divergent growth patterns between axolotl development and regeneration (Fig. 5**b**). CellSTIC further showed that the corresponding communication networks did not follow a single maturation trajectory, but instead encoded organ-specific and process-specific remodelling programs.

**Fig. 5.**
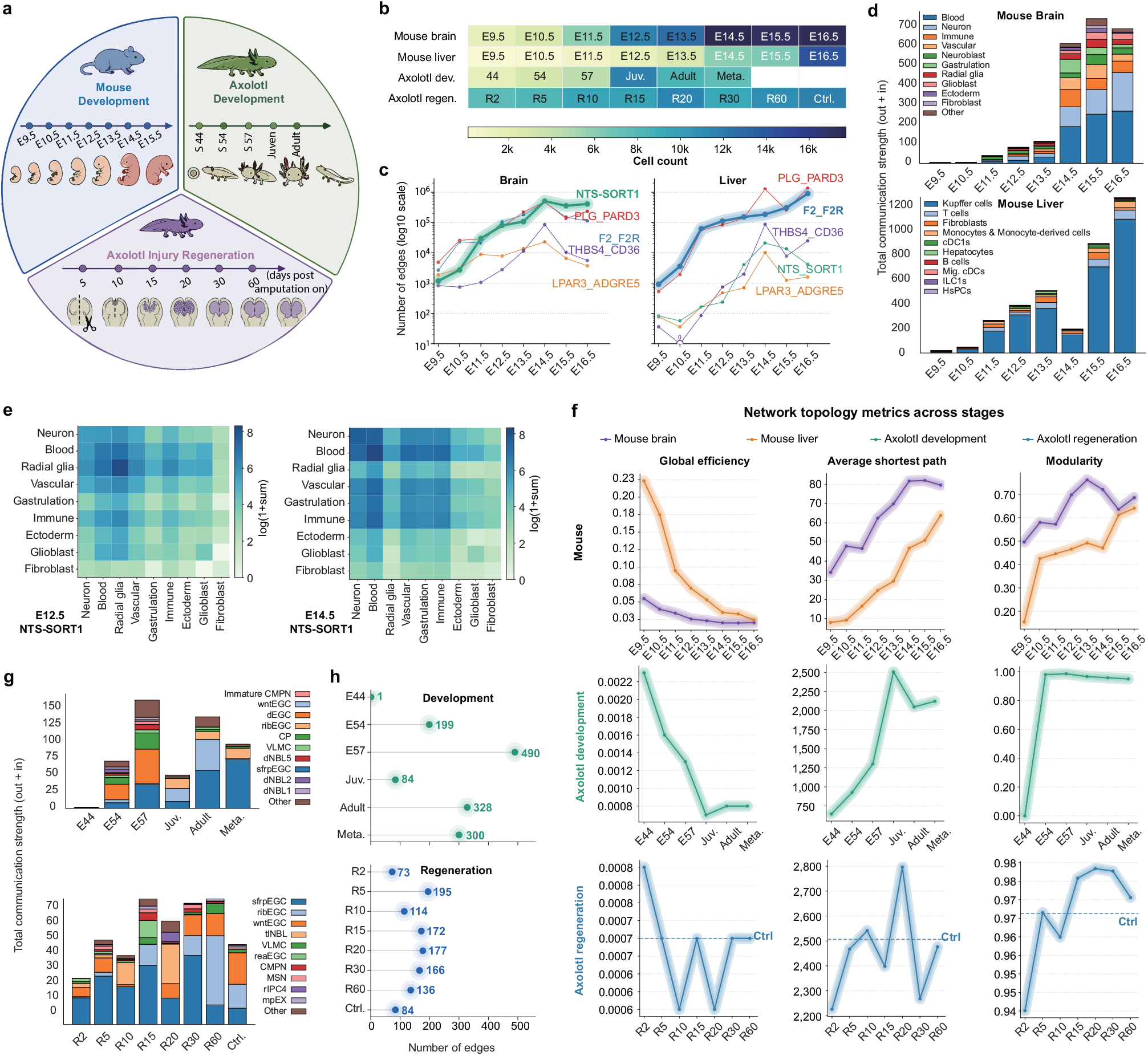
Comparative cell–cell communication dynamics during mouse embryogenesis and axolotl telencephalon development and regeneration. **a**, Schematic overview of the analysed systems and sampling schemes, including mouse embryogenesis, axolotl telencephalon development, and axolotl telencephalon regeneration. **b**, Changes in cell number across stages, including mouse brain and liver during embryogenesis and axolotl telencephalon during development and regeneration. **c**, Changes in the number of network edges for five representative ligand–receptor interaction networks across mouse embryonic stages. **d**, Major cell-type contributors to total communication strength (outgoing plus incoming) in the Brain NTS-SORT1 and Liver F2-F2R communication networks across mouse embryogenesis; colors denote contributing cell types. **e**, Cell-type communication matrices for the Brain NTS-SORT1 communication network at E12.5 (left) and E14.5 (right); color intensity indicates communication strength. **f**, Stage-dependent changes in global network properties of the Brain NTS-SORT1 and Liver F2-F2R communication networks, and of the WNT7B-FZD5 communication network during axolotl telencephalon development and regeneration, including global efficiency, average shortest path length, and modularity. **g**, Major cell-type contributors to total communication strength (outgoing plus incoming) in axolotl telencephalon communication networks during development (top) and regeneration (bottom); colors denote contributing cell types. **h**, Changes in the number of network edges in the WNT7B-FZD5 communication network during axolotl telencephalon development (top) and regeneration (bottom).

In mouse embryogenesis, CellSTIC resolved two distinct developmental logics. In the brain, the NTS-SORT1 network became prominent after E13.5, whereas in the liver, F2-F2R remained the dominant communication axis across mid-to-late development (Fig. 5**c** and Extended Data Fig. 5**b**). Both networks showed decreasing global efficiency together with increasing average shortest path length and modularity, indicating a transition from early centralized communication towards later community-structured organization (Fig. 5**f**). However, the biological implementation of this transition differed between organs. Brain communication became less dominated by a small number of high-connectivity nodes and was progressively organized around emerging neural compartments, while retaining long-tailed degree and strength distributions. Spatially, brain communication decayed gradually with distance, consistent with broader neural-field organization. By contrast, liver communication showed stronger short-range dependence and remained centred on macrophage-associated interactions, consistent with a fetal liver niche in which macrophages contribute to erythroid maturation and haematopoietic organization [35] (Figs. 5**d, e** and Extended Data Figs. 5**c**–**e**). This organ-specific divergence was particularly evident in the brain NTS-SORT1 network. At E12.5, communication was dominated by blood, vascular and immature neural-like populations, whereas by E14.5 the same axis was redistributed towards fore-brain GABAergic neurons, glutamatergic neuroblasts and radial glia (Fig. 5**e**). This shift indicates qualitative rewiring of sender-receiver relationships rather than a simple increase in edge number, and is consistent with the role of SORT1 in neuronal lipid uptake and metabolic substrate utilization [36].

The axolotl telencephalon further tested whether CellSTIC could distinguish developmental maturation from injury-induced reconstruction within the same tissue. WNT7B-FZD5 was reconstructed in both developmental and regenerative series, but showed distinct quantitative and topological behaviour (Figs. 5**f**–**h** and Extended Data Figs. 6**c**–**f**). During development, WNT7B-FZD5 communication shifted from an early, sparse state at E44 to a broadly established network from E54 to E57 onward, with increased edge number, reduced global efficiency, longer shortest path length, and higher modularity. These changes indicate the incorporation of WNT7B-FZD5 signalling into a more spatially extended and compartmentalized telencephalon communication network. Regeneration showed a different pattern. WNT7B-FZD5 communication was rapidly redeployed after injury, with fluctuating total strength and cellular contributors across R2 to R60. Edge number showed an early peak at R5, followed by a drop at R10 and later fluctuations, whereas shortest path length and modularity showed transient deviations before partial architectural re-stabilization (Figs. 5**f**–**h**). These results support a disruption-reorganization model in which injury rapidly reallocates WNT7B-FZD5 communication, followed by staged topological adjustment. Consistent with studies showing that axolotl brain regeneration involves injury-induced progenitor-like states and regenerative neurogenesis [37], CellSTIC distinguishes regenerative remodelling from a direct replay of development.

## 3 Discussion

CellSTIC provides a framework for viewing cell–cell communication in spatial multi-omics as an organized feature of tissue architecture rather than as a collection of disconnected ligand–receptor pairs. A central premise of the method is that communication is most informative when resolved across multiple levels of biological organization: from individual molecular interactions to higher-order programs that remain traceable to their underlying evidence. By combining multimodal integration within local tissue neighborhoods with hierarchical semantic organization of ligand–receptor interactions, CellSTIC links molecular evidence, spatial context, and biological interpretation within a unified analytical representation.

Across benchmarks and tissue settings, this representation revealed communication structure along complementary semantic, spatial, and temporal axes. In this sense, the principal contribution of CellSTIC is not only greater robustness of inference, but a shift in how communication can be analysed: as a system that can be compared across contexts, localized within tissue architecture, and interpreted across scales. In immune tissue, this enabled communication to be summarized as branch-restricted, multiscale programs rather than as fragmented interaction candidates. In the brain, it resolved anatomically coherent signaling domains and their local drivers. Across mouse embryogenesis and axolotl telencephalon development and regeneration, it captured coordinated remodeling of communication networks while distinguishing both organ-specific and process-specific trajectories. Together, these findings suggest that structured representations may offer a more general basis for interpreting intercellular signaling in complex spatial systems.

The choice of hierarchy construction should be guided by the balance between structural regularity and biological interpretability required for a given analysis. Balanced hierarchies are most useful when training stability and robustness are the primary concern, particularly in the presence of strong class imbalance. LLM-guided hierarchies provide an intermediate solution by preserving approximate structural balance while improving semantic coherence. By contrast, biologically informed hierarchies are most appropriate when traceable functional meaning is prioritized over formal regularity. These strategies are therefore best viewed as complementary modes of organizing the same communication space, each emphasizing a different trade-off between optimization, semantic structure, and prior biological knowledge.

This framework may also be useful experimentally because it allows communication to be interrogated at multiple levels of resolution. Broad modules can first be used to identify dominant signaling themes, and these can then be refined into higher-priority ligand–receptor pairs and spatially proximal cell-pair interactions for targeted follow-up. Such a progression may be especially valuable in complex tissues, where communication is spatially heterogeneous and mechanistically layered. The comparative analyses presented here further suggest that this multilevel representation is useful not only for describing stable tissue organization but also for distinguishing different modes of temporal remodeling, including developmental consolidation and regeneration-associated network reassembly. More broadly, CellSTIC provides a generalizable foundation for linking tissue organization to intercellular signaling in situ. As spatial data quality, molecular annotations, and prior knowledge continue to improve, frameworks of this kind should further expand the ability to generate mechanistic and experimentally testable hypotheses from spatial multi-omic data.

## 4 Methods

### 4.1 Datasets

#### Spatial Multimodal Dataset

To evaluate the model’s ability to identify spatial regions in spatial multi-omics data, we used the same synthetic spatial multimodal dataset as SpatialGlue [18], generated following Townes et al. [22]. Spatial structure is constructed using the ggblocks model with non-negative spatial decomposition, while three modalities are simulated: RNA for gene expression, ADT (antibody-derived tags) for protein abundance, and ATAC (assay for transposase-accessible chromatin) for chromatin accessibility. The dataset contains 1,296 cells with spatial-domain annotations. RNA is a gene expression matrix sampled from a zero-inflated negative binomial distribution with four independent factors, and ADT is a protein expression matrix sampled from a negative binomial distribution with four independent factors. ATAC is generated using the dataset’s original setting and used for downstream modeling.

#### CCC Multimodal Dataset

We used a multimodal synthetic dataset generated by scMultiSim [21] to evaluate cell–cell communication prediction. scMultiSim simulates “true” intracellular expression using a kinetic model and then adds technical noise to mimic real data characteristics. Biological structure is encoded by latent variables, including cell identity factors (CIFs), gene identity vectors (GIVs), and region identity vectors (RIVs). The dataset contains RNA and ATAC modalities with spatial coordinates and provides ground-truth CCC annotations. We generated eight replicate samples (four cell types per sample) with 120 genes, and enabled six ligand–receptor pairs for CCC evaluation.

#### Human lymph node dataset

We used the human lymph node spatial multi-omics dataset from SpatialGlue, which contains spatial transcriptomics and joint RNA-protein measurements. We analysed region A1 (3,484 spots; 18,085 genes; 31 proteins). Mitochondrial genes were removed, and genes expressed in fewer than 50 spots were filtered out. Modality-specific dimensionality reduction was performed: for RNA, 3,000 HVGs were selected, and 500 PCs were retained; for ADT, data were normalized, and 30 PCs were retained. As cell-type labels are not provided, we applied CellTypist for automatic annotation.

#### Mouse brain 5M dataset

We used the Mouse Brain 5M dataset from Guo et al. [38] (Spatial-Mux-seq), which profiles ATAC, H3K27me3, H3K27ac, proteins, and RNA in mouse brain tissue and integrates spatial ATAC-RNA-seq with CUT&Tag-RNA-seq. The dataset contains 10,000 cells/pixels (48,440 genes; 16,835 H3K27ac peaks; 6,979 H3K27me3 peaks; 21,091 ATAC peaks; 131 proteins). Quality control removed pixels with < 200 detected genes and genes expressed in < 200 pixels. RNA counts were library-size normalized and log-transformed (Scanpy), followed by selecting 3,000 HVGs and PCA (top 50 PCs). Peak/chromatin data were embedded with LSI into 50 dimensions aligned to the RNA embedding space. Cell-type labels were obtained from the online portal (https://brain-map.org/bkp/browse) and used downstream.

#### Human skin and tonsil datasets

We used the human skin and tonsil spatial CITE-seq datasets from Liu et al. [39], enabling co-localized measurement of multiplexed proteins and whole-transcriptome profiles. The skin dataset contains 15,486 genes and 283 proteins, and the tonsil dataset contains 28,417 genes and 283 proteins. Mitochondrial genes were removed, and a unified pipeline was applied: for RNA, 3,000 HVGs were selected, followed by normalization, log transformation, and PCA (250 PCs); for ADT, CLR normalization was applied, and PCA retained 250 PCs.

#### Mouse embryonic dataset

Mouse embryonic spatial transcriptomic data were obtained from MOSTA (Stereo-seq), comprising 53 sagittal sections across eight stages (E9.5–E16.5), with cell counts ranging from 5,913 (E9.5) to 121,767 (E16.5) [33]. RNA preprocessing removed mitochondrial genes, filtered genes expressed in fewer than 100 cells, selected 3,000 HVGs (Seurat), normalized counts to 1,000, applied log1p transformation, and performed PCA (500 PCs). Cell types were annotated using the CellTypist tool for the mouse developing brain tissue.

#### Axolotl telencephalon dataset

Axolotl telencephalon spatial transcriptomic data were obtained from ARTISTA (Stereo-seq), a public atlas of axolotl brain development and regeneration at single-cell spatial resolution [34]. The dataset covers six developmental stages (Stage44, Stage54, Stage57, juvenile control, adult, and metamorphosed) and seven regeneration time points (2, 5, 10, 15, 20, 30, and 60 DPI). In our study, a raw copy of the expression matrix was retained before preprocessing. Mitochondrial genes were annotated on the basis of gene names and used for quality-control metric calculation. Highly variable genes were then identified using the Seurat v3 method with the top 3,000 genes retained. Counts were normalized to a total of 1,000 per cell and log-transformed using log1p for downstream analysis.

### 4.2 CellSTIC Overview

CellSTIC (Fig. 1) is a spatial multi-omics framework for inferring cell–cell communication. Taking spatial coordinates and multimodal measurements as input, the Multimodal Evidence Graph Constructor builds a spatially constrained communication backbone graph and strengthens its structure by integrating modality-specific evidence from local neighborhoods and global clustering patterns. The Multimodal Evidence Integrator then learns cross-modal fused representations, jointly guided by a reconstruction objective and a functional region-oriented objective to stabilize alignment and capture spatial domain organization. To improve interpretability, the Ligand–Receptor Semantic Tree Builder organizes ligand–receptor interaction types into a hierarchical semantic tree, enabling multi-level functional analysis of communication patterns. Finally, the Cell–Cell Communication Predictor performs hierarchical training with a self-supervised edge masking-reconstruction scheme and outputs communication strengths and interaction-type predictions at multiple semantic resolutions. In addition, CellSTIC explicitly encodes edge semantics (distance, directionality, and interaction strength) and employs the Hyperplane Orthogonal Decomposition Graph Neural Network for node-edge co-updating, thereby improving the modeling of spatially dependent communication mechanisms.

### 4.3 Communication Graph Generation Based on Spatial Constraints and Multimodal Evidence

Cell–cell communication is mediated by ligand–receptor interactions and is inherently spatial, driven by proximity and supported by multimodal signals. We propose the Multimodal Evidence Graph Constructor (MEGC; Extended Data Fig. 1**a**) to build a communication graph under the principle of spatial constraints and multi-modal evidence. Using a ligand–receptor database, it constructs a typed, directed backbone graph constrained by spatial distance and ligand–receptor strength. It further computes intercellular similarity within each modality and encodes these similarities together with spatial coordinates and directionality as edge features, producing a mechanism-consistent input for downstream modeling.

#### 4.3.1 Feature-Driven Local-Global Modality Graphs

To incorporate structural information from multimodal measurements, the Multimodal Evidence Graph Constructor builds a feature graph for each modality. Given *N*_spot_ spots and *N*_modal_ modalities, modality *m* is represented by a feature matrix 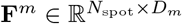, where the *u*-th row 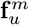 is the feature vector of spot *c*_*u*_. For each modality *m*, we construct two graphs on the shared node set 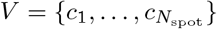:

1. a KNN graph 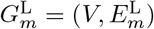, which captures local neighborhood structure by linking each node to its top-*K* nearest neighbors in the modality feature space using cosine similarity;
2. a clustering graph 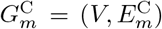, which encodes global population structure by fully connecting nodes within the same unsupervised cluster. To mitigate topological imbalance caused by large clusters, we iteratively split oversized clusters into a predefined size range following Hou et al. [40].

We then define the modality graph *G*^*m*^ = (*V, E*^*m*^, **X**^*m*^, ℰ^*m*^) to integrate local and global information, where 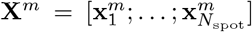 with 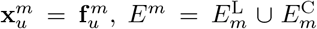, and 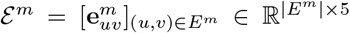. For each 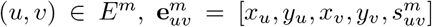, where (*x*_*u*_, *y*_*u*_) and (*x*_*v*_, *y*_*v*_) are spatial coordinates and 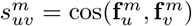 is the modality-specific similarity.

### 4.3 LR-Constrained Spatial Communication Graph

Given *N*_LR_ ligand–receptor pairs 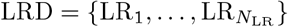 and RNA data 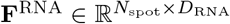, we define a typed, directed backbone graph *G*_*B*_ = (*V, E*_*B*_, **X**_*B*_, ℰ_*B*_), where 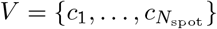 and 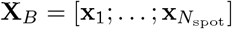 with **x**_*u*_ the *u*-th row of **F**^RNA^. Each directed typed edge is represented as a triplet (*u, v, ℓ*) ∈ *E*_*B*_, indicating a potential communication from node *u* to node *v* associated with ligand–receptor pair LR_*ℓ*_. We further define the edge-feature matrix as 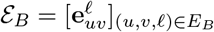.

For each ordered spot pair (*u, v*) and ligand–receptor pair LR_*ℓ*_, we construct a directed edge (*u, v, ℓ*) ∈*E*_*B*_ when both spatial proximity and expression intensity criteria are satisfied. Specifically, the spatial distance between spots *u* and 𝒱 is required to satisfy 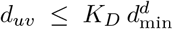, where 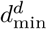 denotes the minimum distance between adjacent spots, and *K*_*D*_ defines the maximum interaction range as a multiple of this minimum spot spacing. The expression criterion requires that the ligand expression at the source spot and the receptor expression at the target spot exceed ligand- and receptor-specific intensity thresholds, 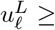 Quantile(*L*^*ℓ*^, *P*_*I*_) and 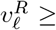 Quantile(*R*^*ℓ*^, *P*_*I*_). Here, *L*^*ℓ*^ and *R*^*ℓ*^ denote the expression vectors of ligand *ℓ* and receptor *ℓ* across all spots, respectively. Accordingly, Quantile(*L*^*ℓ*^, *P*_*I*_) and Quantile(*R*^*ℓ*^, *P*_*I*_) represent the *P*_*I*_-th quantiles of the ligand and receptor expression distributions across all spots. The parameter *P*_*I*_ therefore controls the stringency of the expression intensity threshold. Multiple ligand–receptor pairs may induce parallel edges between the same (*u, v*). For each edge (*u, v, ℓ*), the edge feature is 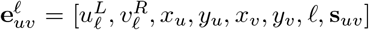, where (*x*_*u*_, *y*_*u*_) and (*x*_v_, *y*_v_) are spatial coordinates and 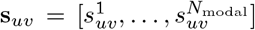 collects cross-modality similarities. Each 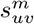 is taken from the modality graph *G*^*m*^ as the similarity term on edge (*u, v*) when (*u, v*) *E*^*m*^. Parallel edges share the same coordinates and **s**_*uv*_, while 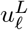 and 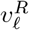 vary with LR_*ℓ*_, preserving evidence for distinct ligand–receptor mechanisms under the same spatial constraint.

### 4.4 Semantics Tree Construction for Structured Knowledge Integration

Ligand–receptor interaction types are often semantically related and highly imbalanced, where a few frequent pairs dominate training. In addition, a flat list of ligand–receptor pairs cannot represent communication functions at multiple biological scales. We therefore propose the Ligand–Receptor Semantics Tree Builder (LRSTB), which constructs a ligand–receptor semantics tree and supports three optional strategies: a balanced strategy, a large language model-guided strategy, and a biological prior strategy. The resulting hierarchy groups ligand– receptor pairs into interpretable functional modules and converts edge-type prediction into a coarse-to-fine layered classification problem, which explicitly encodes label relatedness and mitigates long-tail effects.

#### 4.4.1 Definition of the LRST Structure

Given a ligand–receptor list 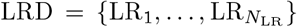, we define the ligand–receptor semantics tree as a hierarchy of height *H*,

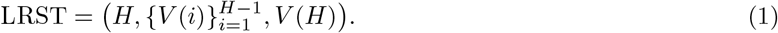

Here, *V* (*i*) is the set of virtual nodes at level *i* for *i* = 1, …, *H*™1, and the leaf level *V* (*H*) is in one-to-one correspondence with LRD, where each leaf represents one ligand–receptor pair. Level 1 contains a single root node, and each internal node represents the subset of ligand–receptor pairs covered by its subtree. Let *n*_*i*_ = |*V* (*i*)| be the number of nodes at level *i*. For node *k* at level *i*, denoted node 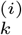, its children are 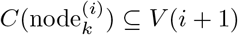, and its covered ligand–receptor set is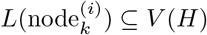.

LRSTB supports three optional construction strategies that share the same input LRD and output the tree structure with the ligand–receptor subsets for internal nodes: a balanced strategy (BAS), a large language model-guided strategy (LLMS), and a biological prior strategy (BIOS) (Extended Data Fig. 1**b**).

#### 4.4.2 Balanced Strategy

The balanced strategy does not use biological priors. Given *N*_LR_, it builds an *N*-ary tree that is as balanced as possible to reduce class size discrepancies across branches during hierarchical training. The tree height *H* is set by a piecewise rule:

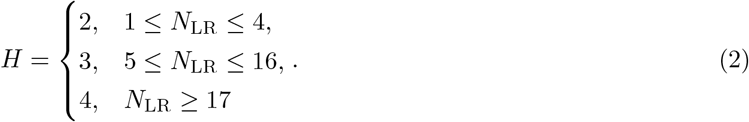

Level 1 is a single root node. When *H* = 2, the leaf level directly contains the *N*_LR_ ligand–receptor nodes. When *H* ≥ 3, the strategy chooses per-level branching factors by approximate integer factorization so that the number of leaves under nodes at the same level is as uniform as possible. For *H* = 3, it selects *n*_2_ = *X* and assigns about *Y* leaves per level-2 node such that *X*·*Y* ≈*N*_LR_, distributing any remainder across a few branches so that the size difference is at most 1. The case *H* = 4 is handled similarly by choosing *X* ·*Y*· *Z*≈ *N*_LR_ and spreading the non-divisible part across last-level branches with at most a one-leaf difference.

### 4.4.3 LLM-Guided Strategy

The large language model-guided strategy uses the balanced strategy to fix the tree height *H* and the number of nodes at each level, and then builds a semantically coherent hierarchy while keeping within-level sizes approximately balanced. It consists of four steps.

#### Semantic embedding

For each ligand–receptor pair LR_*ℓ*_, we form a text description from CellChat v2 annotations and compute an embedding ***θ***_*ℓ*_ = Φ(LR_*ℓ*_). We also embed each virtual-node topic for later matching.

#### Virtual node generation

Starting from level 2, for each parent node we prompt a large language model with the parent topic and its covered ligand–receptor texts to generate the required number of child topics, constrained to be distinct within the same level. Each topic is then embedded.

#### Matching with verification

We first select candidates by cosine similarity between topic and ligand–receptor embeddings, keeping pairs with similarity at least *T*_*s*_. We then use batched large language model verification to filter candidates with weak semantic or functional relatedness.

#### Lightweight balancing

We make small adjustments so that node sizes within the same level differ by at most one, refining candidates near *T*_*s*_ and, when needed, truncating or supplementing using the large language model ranking. Any remaining unmatched ligand–receptor pairs are assigned to the most similar topic.

### 4.4 Biological Prior Strategy

The biological prior strategy prioritizes interpretability by using biologically meaningful CellChat v2 annotations to progressively group LR interactions into a hierarchy with traceable semantics. We apply a four-level rule set: the root filters candidates by interaction validity; the next level groups interactions by functional annotation and whether they are neurotransmitter-related; the third level refines groups using ligand and receptor secretion or localization attributes such as secreted type; and the final level corresponds to individual ligand–receptor pairs. Because biological semantics are prioritized, branch sizes are not necessarily balanced.

### 4.5 Multimodal Evidence Integration in Unified Latent Space

Cell–cell communication is supported by multiple sources of evidence, including molecular states, spatial neighborhoods, and functional phenotypes. Because any single modality captures only part of this evidence, inference can be fragile to noise, scale mismatch, or missing measurements. We therefore propose the Multimodal Evidence Integrator, which performs within-modality purification, cross-modality alignment, and fusion to project modalities into a unified latent space that strengthens consistent signals and suppresses noisy or conflicting ones, yielding robust representations for spatial domain identification and downstream cell–cell communication prediction.

For modality ^*m*^ with feature matrix 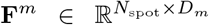, we construct its modality graph *G*^*m*^ = (*V, E*^*m*^, **X**^*m*^, ℰ^*m*^) using the Multimodal Evidence Graph Constructor and obtain a low-dimensional embedding with a modality-specific graph encoder,

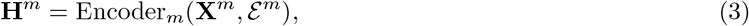

where 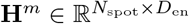 and the encoder stacks multiple Hyperplane Orthogonal Decomposition Graph Neural Network layers to jointly update node and edge features. To preserve modality-specific information, we reconstruct the input with a modality decoder implemented as a multilayer perceptron,

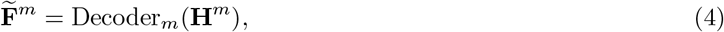

where 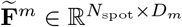. After processing all modalities, we concatenate their embeddings and fuse them with a multilayer perceptron fusion block,

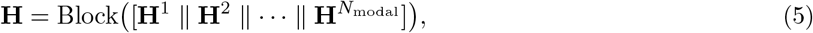

where 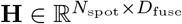. The fused feature vector 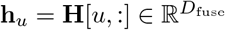 serves as the initial latent representation of spot *u*. This representation is further optimized in the subsequent tree-guided self-supervised CCC learning stage; for simplicity, we continue to denote the refined embedding as **h**_*u*_. The final **h**_*u*_ is then used for functional spatial domain clustering and downstream cell–cell communication prediction.

### 4.6 Tree-Guided Self-Supervised CCC Prediction

CCC graphs typically lack reliable edge labels, making supervised training impractical. We propose a CCC predictor trained on the backbone graph *G*_*B*_ = (*V, E*_*B*_, **X**_*B*_, ℰ_*B*_) using a self-supervised edge masking and reconstruction objective. The model uses an encoder with two decoders: a structure decoder to recover masked edges and a type decoder to predict interaction types. Training follows the ligand–receptor semantics tree in a top-down manner. At each level, ligand–receptor labels are mapped to that level’s virtual types, and the model jointly reconstructs masked edges to capture communication topology and predicts virtual types to learn semantically discriminative edge representations across levels.

#### 4.6.1 Hierarchical Edge-Type Mapping

Consider the backbone graph *G*_*B*_ = (*V, E*_*B*_, **X**_*B*_, *ε*_*B*_), where each ligand–receptor pair 𝓁 induces a directed edge (*u, v*, 𝓁) ∈ *E*_*B*_ with edge feature 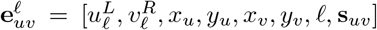. For level *k*, let 𝒱_*k*_ be the set of virtual types and 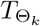 : LRD *→* 𝒱_*k*_ map each ligand–receptor pair to a virtual type. We construct the level-*k* coarsened graph by merging parallel edges that share the same endpoints and virtual type. For each pair (*u, v*) and virtual type *τ*, define the bucket

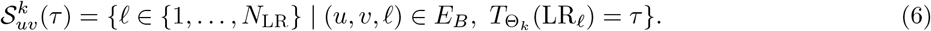

We aggregate only the ligand and receptor intensity terms that vary with 𝓁,

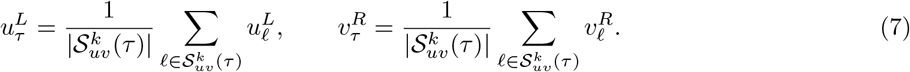

The spatial and similarity terms are shared within each bucket, so they are kept unchanged, equivalently replaced by their within-bucket mean. The coarsened edge feature is 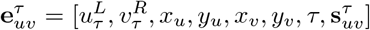, with 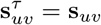 under shared edge evidence. For each level-*k* graph, we mask edges using the same ratio within each virtual type, which reduces dominance by frequent types and improves stability under long-tailed interactions.

#### 4.6.2 Hierarchical Encoder and Decoder

The communication encoder is a stack of Hyperplane Orthogonal Decomposition Graph Neural Network layers that takes node features **X**_*B*_ and level-*k* edge features 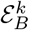 as input and outputs node and edge representations,

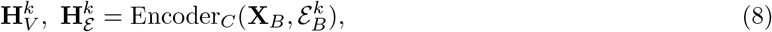

where 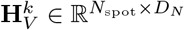 and 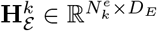, and 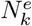 is the number of edges in 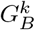. Both the structure decoder and the type decoder use a shared trunk with level-specific heads. The structure decoder predicts the existence probability of a masked edge,

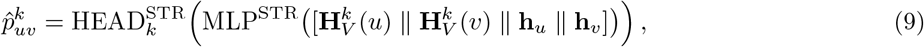

while the type decoder predicts a distribution over level-*k* virtual types,

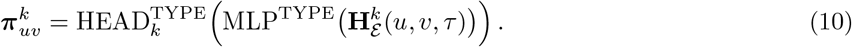

All trunks and heads are implemented as multilayer perceptrons, enabling shared structural learning across levels while maintaining level-specific semantic discrimination for robust cell–cell communication inference.

### 4.7 Hyperplane Orthogonal Decomposition Graph Neural Network

A spot-level communication graph combines sequencing features and spatial geometry. Since most graph neural networks do not explicitly exploit edge features, spatial signals are often merged into node inputs and can be overwhelmed by high-dimensional omics features, weakening spatially dependent communication modeling. We therefore use sequencing measurements as node features and encode spatial cues, including coordinates and ligand–receptor strength, as edge features, and apply the Hyperplane Orthogonal Decomposition Graph Neural Network (HODGNN) to perform edge-guided aggregation in orthogonal subspaces, improving spatially aware communication inference.

#### Input

Given a directed graph *G* = (*V, E*, **X**^(*i−*1)^, *ε*^(*i−*1)^), where 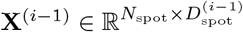 is the node-feature matrix and 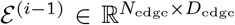 is the edge-feature matrix, the representation of node *u* is 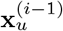 and the representation of edge 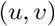 is 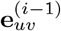.

#### Edge-driven orthogonal subspaces

At layer *i*, we maintain *N*_space_ trainable vectors 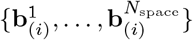 and orthonormalize them to obtain 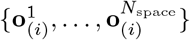 using Gram–Schmidt. Each edge is then projected onto this basis to produce subspace coefficients

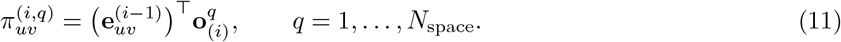

This orthogonal projection disentangles edge semantics into independent subspaces for structured message modulation.

#### Edge-modulated attention aggregation

For each subspace *q*, we compute a node-pair attention score

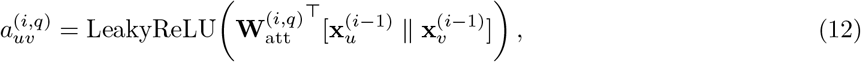

where 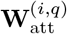 is a learnable attention parameter for subspace *q* at layer *i*. We then normalize the edge-modulated attention weights as

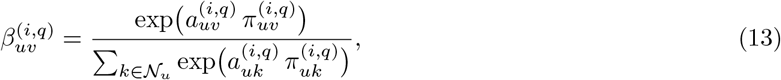

where 𝒩_*u*_ denotes the neighbor set of node *u*. The subspace-specific aggregated node representation is

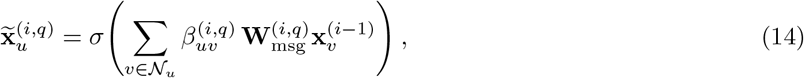

where 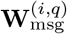 is a learnable message transformation matrix for subspace *q* at layer *i*, and *σ*(·) denotes the activation function. Edge coefficients directly modulate attention, strengthening spatially aware message passing.

#### Cross-subspace fusion and edge update

Because different subspaces correspond to different edge semantic perspectives, layer *i* adaptively fuses 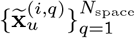 using learned weights to emphasize informative subspaces and suppress noisy ones:

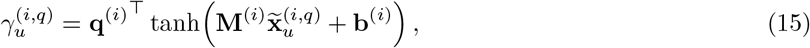

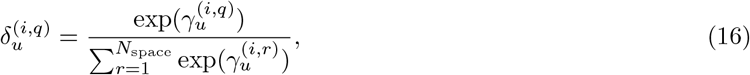

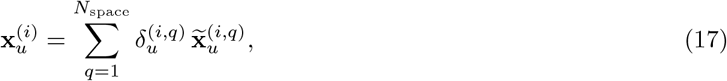

where **q**^(*i*)^ is a learnable scoring vector, and **M**^(*i*)^ and **b**^(*i*)^ are learnable projection parameters. To keep edge-level semantics aligned with node aggregation, we update each edge as a weighted combination of the basis vectors,

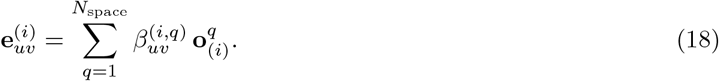

This layer explicitly uses edge features to control message passing through orthogonal subspaces, yielding node and edge representations that better capture spatially dependent communication.

### 4.8 Optimization

We decouple representation learning and spatial domain partitioning from communication network prediction using a two-stage training scheme. In Stage 1, the model learns fused embeddings and derives stable spatial functional domains. In Stage 2, the fusion module is initialized from Stage 1 and jointly optimized with the communication predictor to reconstruct edges and predict their types along the hierarchy.

*Stage 1*. Let *M*_*F*_ denote the modalities used for fusion. The objective combines modality reconstruction and unsupervised clustering:

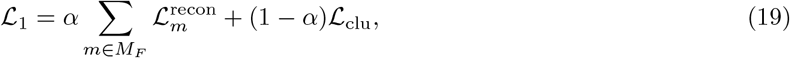

where *α* is a weighting coefficient. For each modality *m*, we use an MSE reconstruction loss

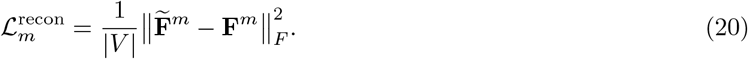

The clustering loss includes a soft *k*-means term with an entropy regularizer. Let **p**_*i*_ = (*p*_*i*,1_, …, *p*_*i,C*_) be the soft assignment of sample *i*:

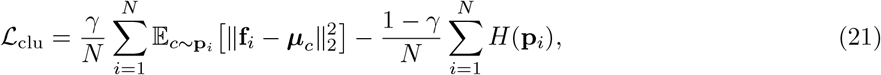

where **f**_*i*_ and ***μ***_*c*_ are 𝓁_2_-normalized embeddings and centroids, *C* is the number of clusters, and 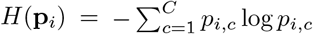. This jointly preserves modality information and yields compact, stable spatial domains.

*Stage 2*. We initialize the fused embeddings and fusion module from Stage 1 and jointly optimize them together with the communication predictor in a top-down manner along the ligand–receptor semantics tree. At level *k*, fine-grained types are mapped to level-*k* virtual types to form the coarsened graph 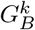. We apply class-balanced masking to obtain masked positive edges 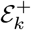 and negative samples 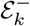, and minimize

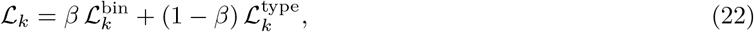

where *β* is a weighting coefficient. The binary reconstruction loss is

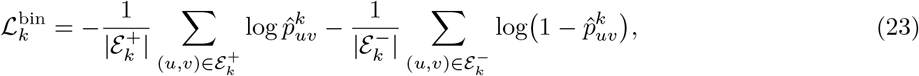

and the type loss is a multiclass cross-entropy over the level-*k* virtual types:

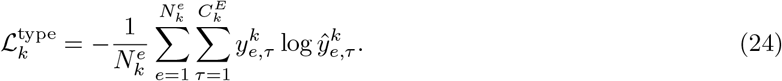

Optimizing 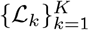 progressively refines communication topology and interaction-type patterns from coarse to fine semantic granularity. By fine-tuning the Stage 1 fusion module together with the communication predictor, coarse-to-fine training couples structure reconstruction with hierarchical type supervision for robust prediction.

## 5 Data Availability

The human lymph node dataset, human skin and tonsil dataset, and mouse brain tissue dataset used in this study were obtained from the Gene Expression Omnibus (GEO) database under accession numbers GSE263617, GSE213264, and GSE263333, respectively. The mouse developmental dataset was collected from the MOSTA database (https://db.cngb.org/stomics/mosta/), and the salamander developmental and injury-repair dataset was downloaded from ARTISTA (https://db.cngb.org/stomics/artista/). The two simulated datasets were derived from the SpatialGlue and scMultiSim studies, respectively, and were downloaded from the public data resources provided in the corresponding papers. In addition, the aggregated raw data are publicly accessible at https://drive.google.com/drive/folders/1vAYlhr8M8tNfU15lU30Z6uv-hj3jhpe7?usp=drive_link.

## 6 Code Availability

An open-source Python implementation of the CellSTIC toolkit, together with scripts for reproducing the analyses, is available at https://github.com/xuyungang/CellSTIC.

## 7 Conflict of Interest

The authors declare that they have no competing interests.

## 8 Acknowledgments

This work was supported by the National Natural Science Foundation of China (62471378, 82541006, and 62171365), the Young Talent Support Plan of Xi’an Jiaotong University (YX6J021), and Shaanxi Province Key Research and Development Projects (2024SF-GJHX-40 and QCYRCXM-2022-209).

---

**Extended Data Fig. 1.**
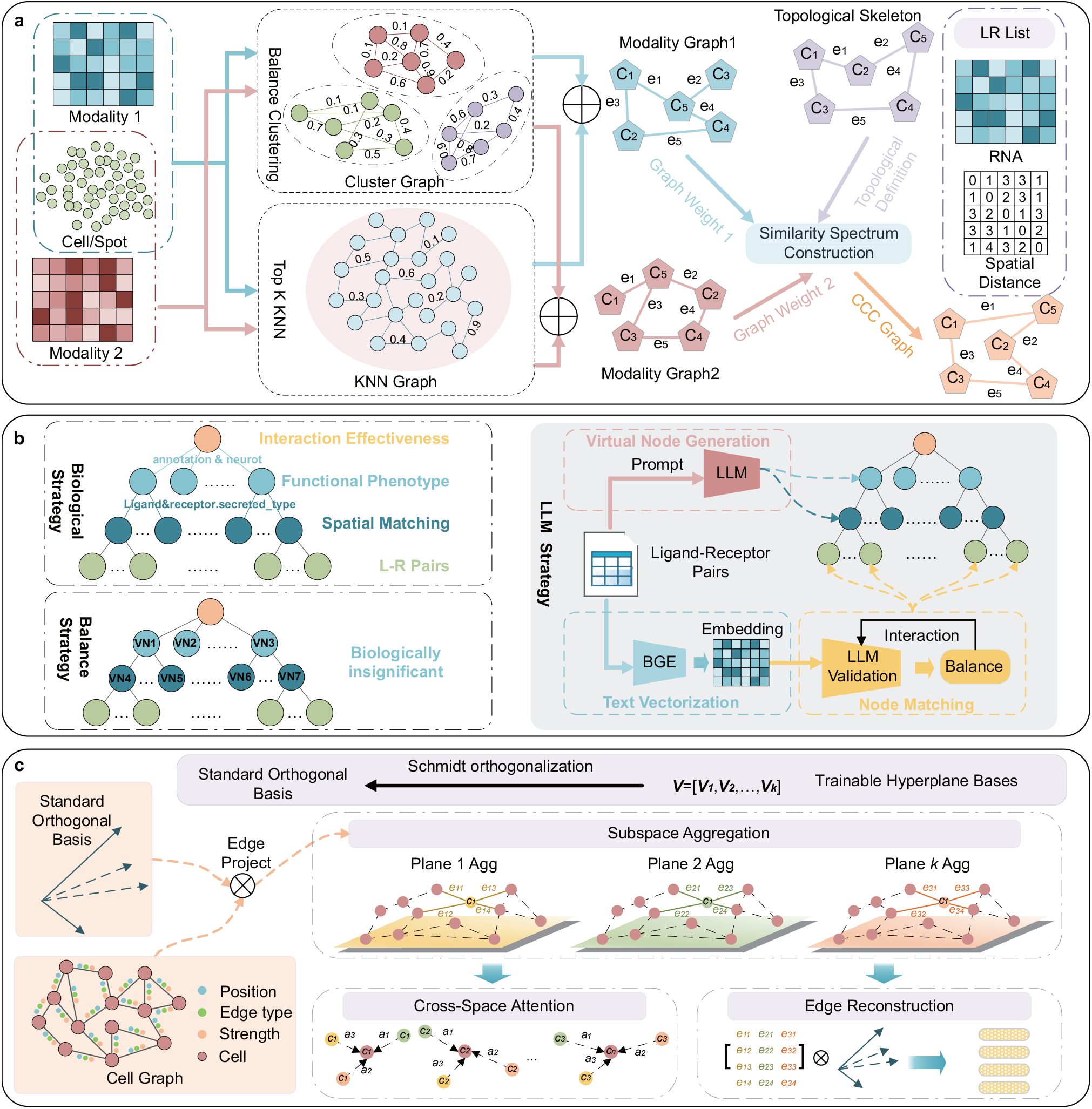
Key components of the CellSTIC framework. **a**, Multimodal Evidence Graph Constructor (MEGC). MEGC constructs modality-specific multi-scale neighbourhoods by integrating clustering structure with *k*-nearest-neighbour graphs, and then refines the spatially constrained communication graph using graph-quality metrics to improve structural consistency and the robustness of cell–cell communication inference. **b**, Ligand–Receptor Semantics Tree Builder (LRSTB). LRSTB builds a hierarchical ligand–receptor semantics tree using complementary balancing, biologically guided, and large-language-model-guided strategies, enabling hierarchical organization of interactions and traceable interpretation from communication modules to individual ligand–receptor pairs. **c**, Hyperplane Orthogonal Decomposition Graph Neural Network (HODGNN). HODGNN models communication graphs with both node and edge features through hyperplane-based orthogonal decomposition and attention-weighted aggregation, enabling joint modeling of node-edge co-occurrence patterns and integration of information across multiple subspaces.

**Extended Data Fig. 2.**
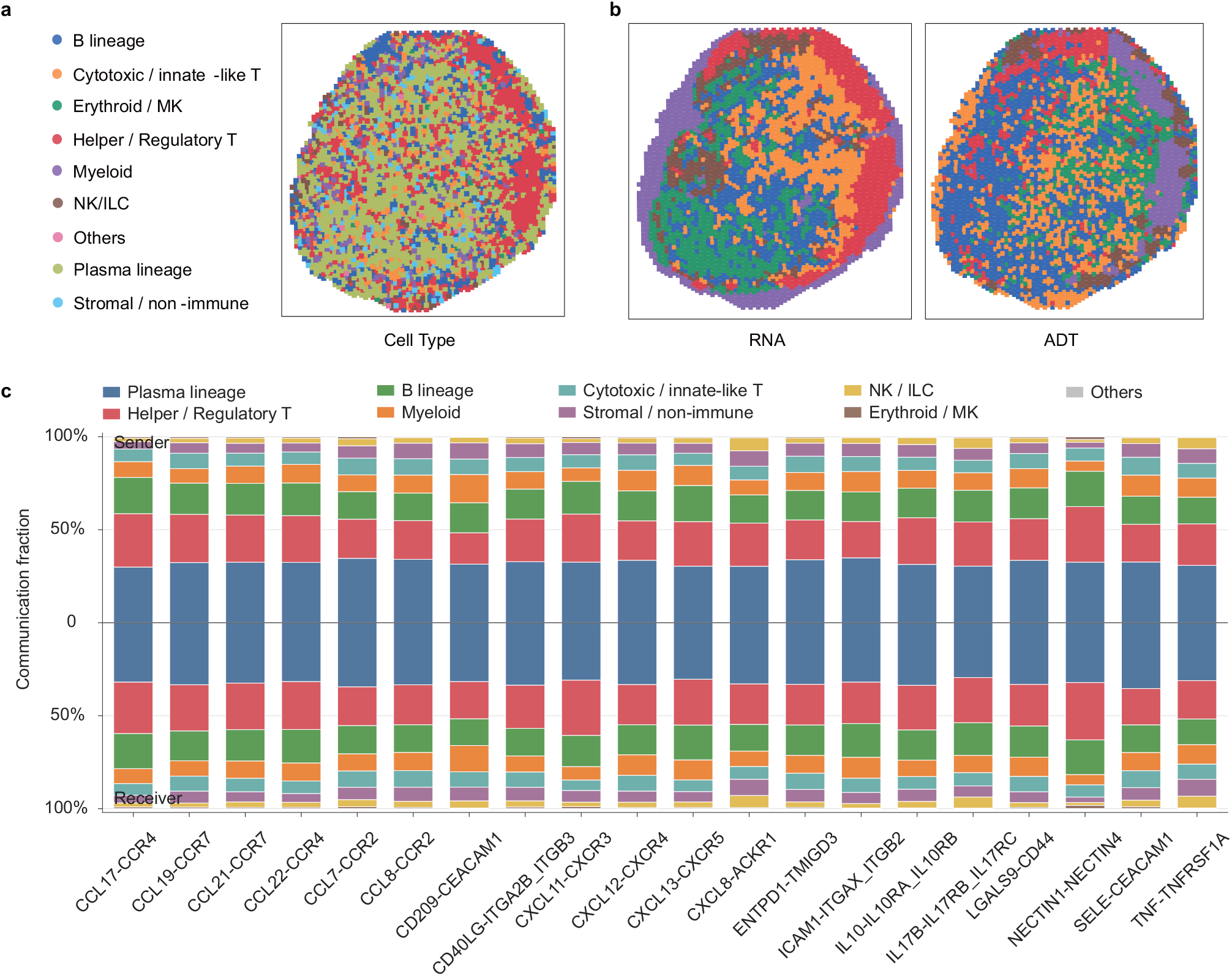
Post hoc cell-type annotations, unimodal clustering, and sender-receiver cell-type composition of representative ligand–receptor pairs in the human lymph node. **a**, Spatial distribution of post hoc cell-type annotations in the human lymph node. These annotations were used for interpretation only and were not provided to CellSTIC during model fitting. **b**, Spatial maps of unimodal clustering based on RNA expression (left) and ADT expression (right). Compared with the communication domains identified by CellSTIC (Fig. 3**a**), these unimodal clustering results capture molecular variation but do not recover the same spatially coherent communication organization. **c**, Sender and receiver cell-type composition of representative ligand– receptor pairs. For each ligand–receptor pair, stacked bars show the fraction of communication contributed by each cell type as sender (top) or receiver (bottom).

**Extended Data Fig. 3.**
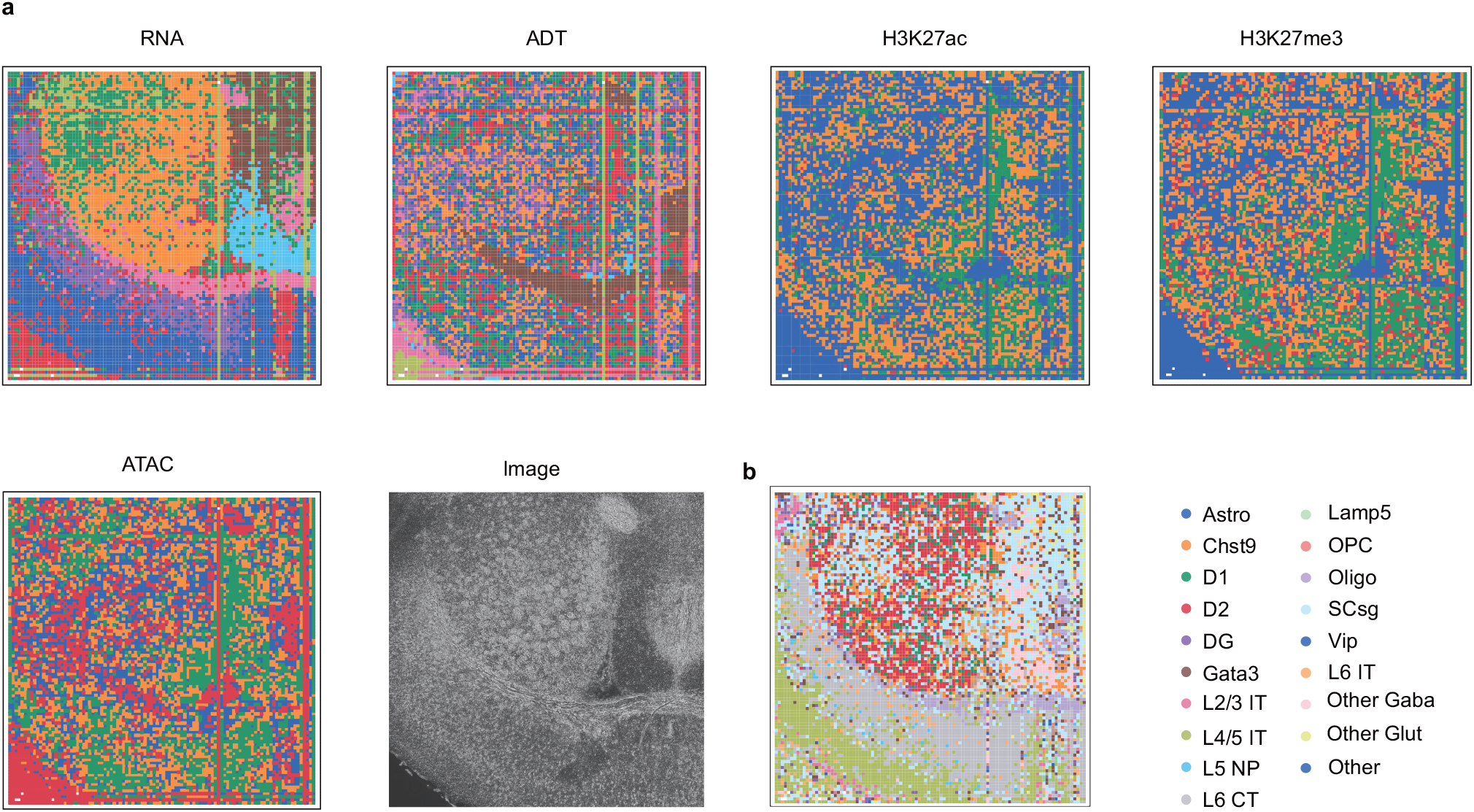
Multimodal spatial domains and cell-type composition in the mouse brain. **a**, Tissue image and spatial partitions inferred from the individual modalities used for integration. **b**, Spatial map of cell-type composition across the same section.

**Extended Data Fig. 4.**
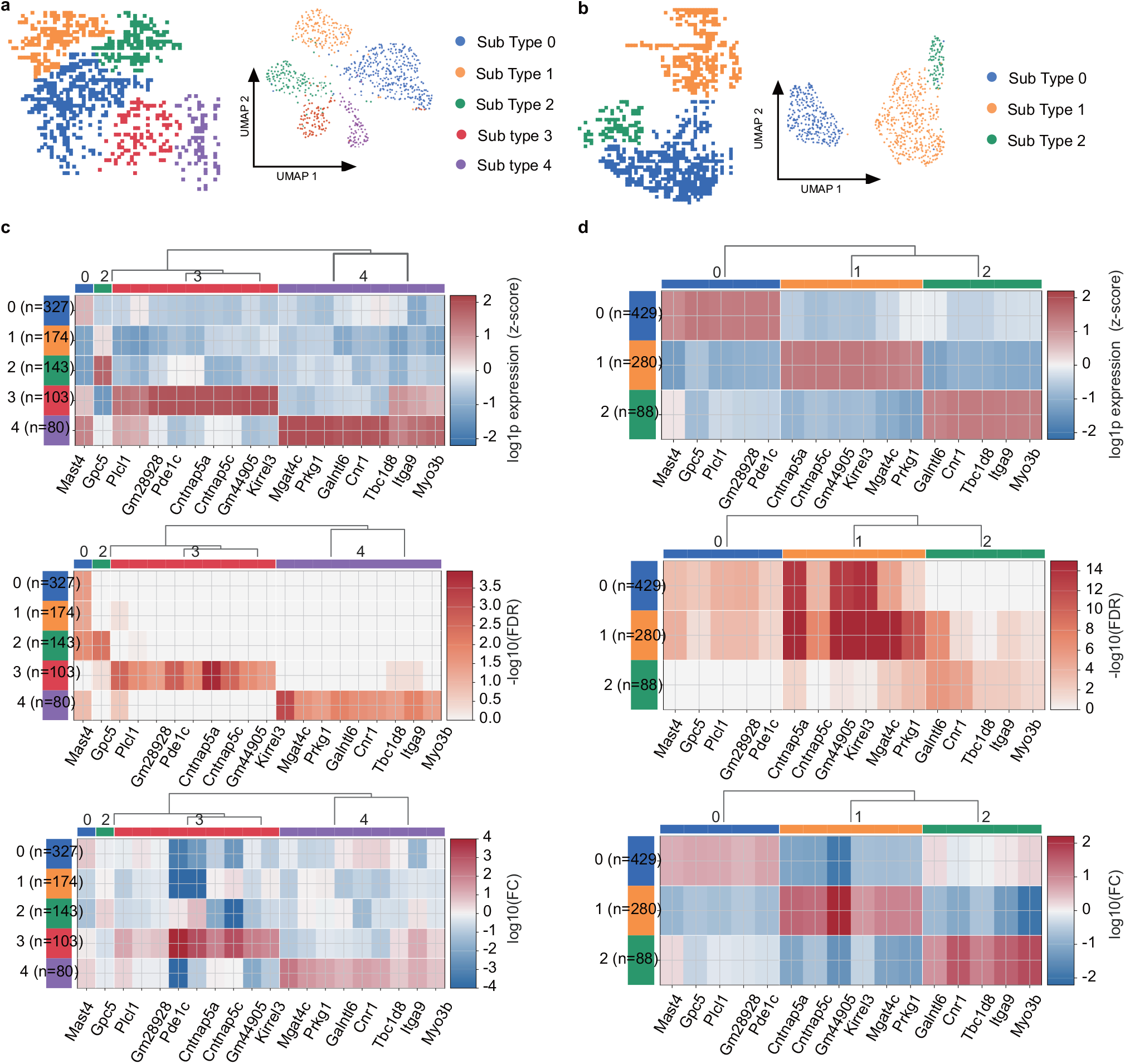
Communication microstructure analysis of selected spatial domains in the mouse brain dataset. **a**,**b**, Spatial maps and UMAP embeddings of detected CCC communities in domains 9 (**a**) and 11 (**b**). **c**,**d**, Differential communication features across communities in domains 9 (**c**) and 11 (**d**), shown by scaled log_2_ expression, *−* log_10_(FDR), and log_2_ fold change.

**Extended Data Fig. 5.**
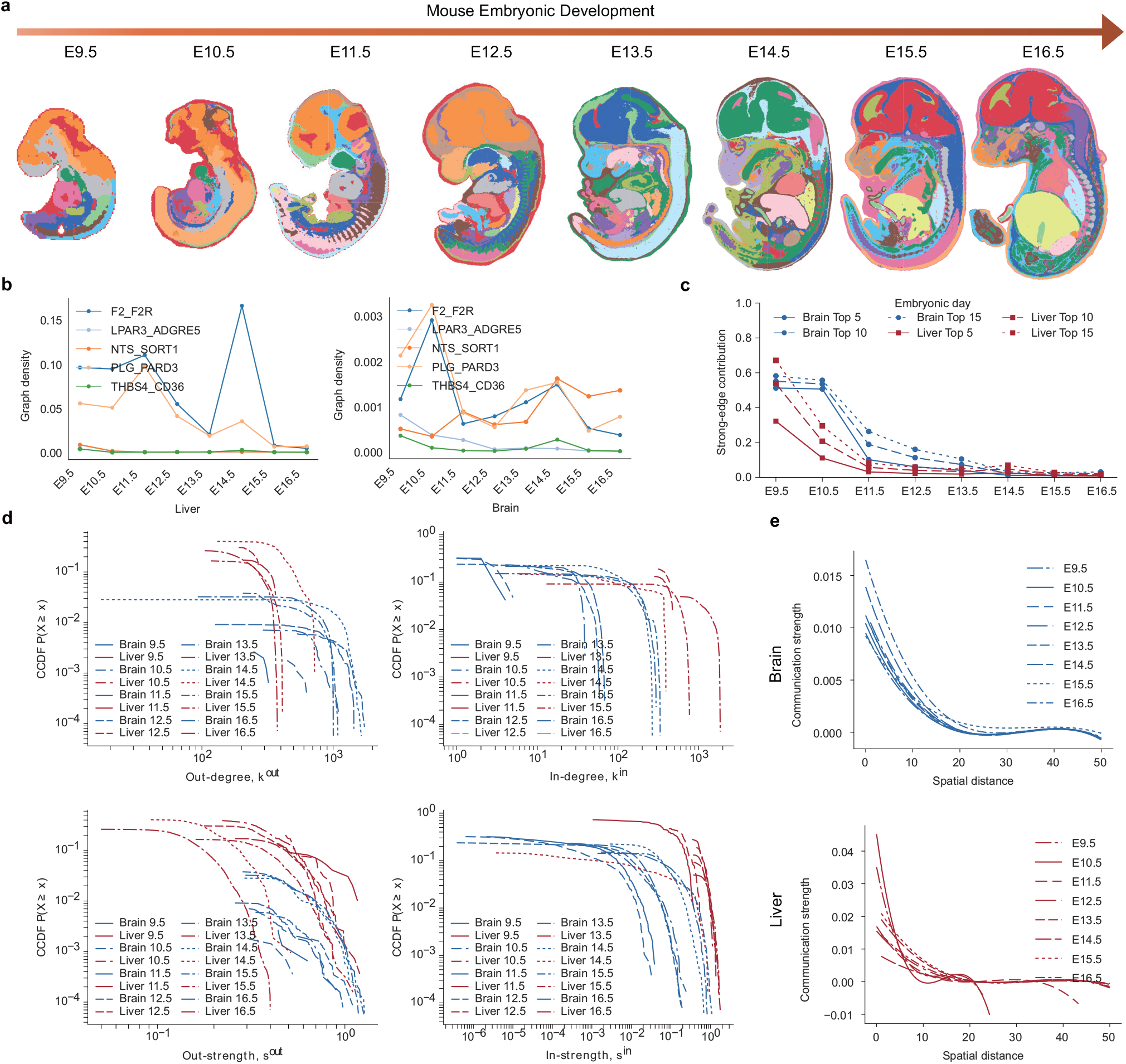
Additional topological characteristics of spatial cell–cell communication networks during mouse embryogenesis. **a**, Overview of the mouse embryonic spatial transcriptomics dataset analysed in this study, showing representative embryo sections across developmental stages. **b**, Graph density of five representative ligand–receptor interaction networks in the brain and liver across mouse embryonic stages. **c**, Fractions of total communication contributed by the most highly connected nodes in the Brain NTS-SORT1 and Liver F2-F2R communication networks, shown for the top 5, top 10, and top 15 nodes. **d**, Complementary cumulative distribution function (CCDF) curves of out-degree, in-degree, out-strength, and in-strength for the Brain NTS-SORT1 and Liver F2-F2R communication networks across developmental stages. **e**, Distance-dependent decay in communication strength for the Brain NTS-SORT1 communication network (top) and the Liver F2-F2R communication network (bottom).

**Extended Data Fig. 6.**
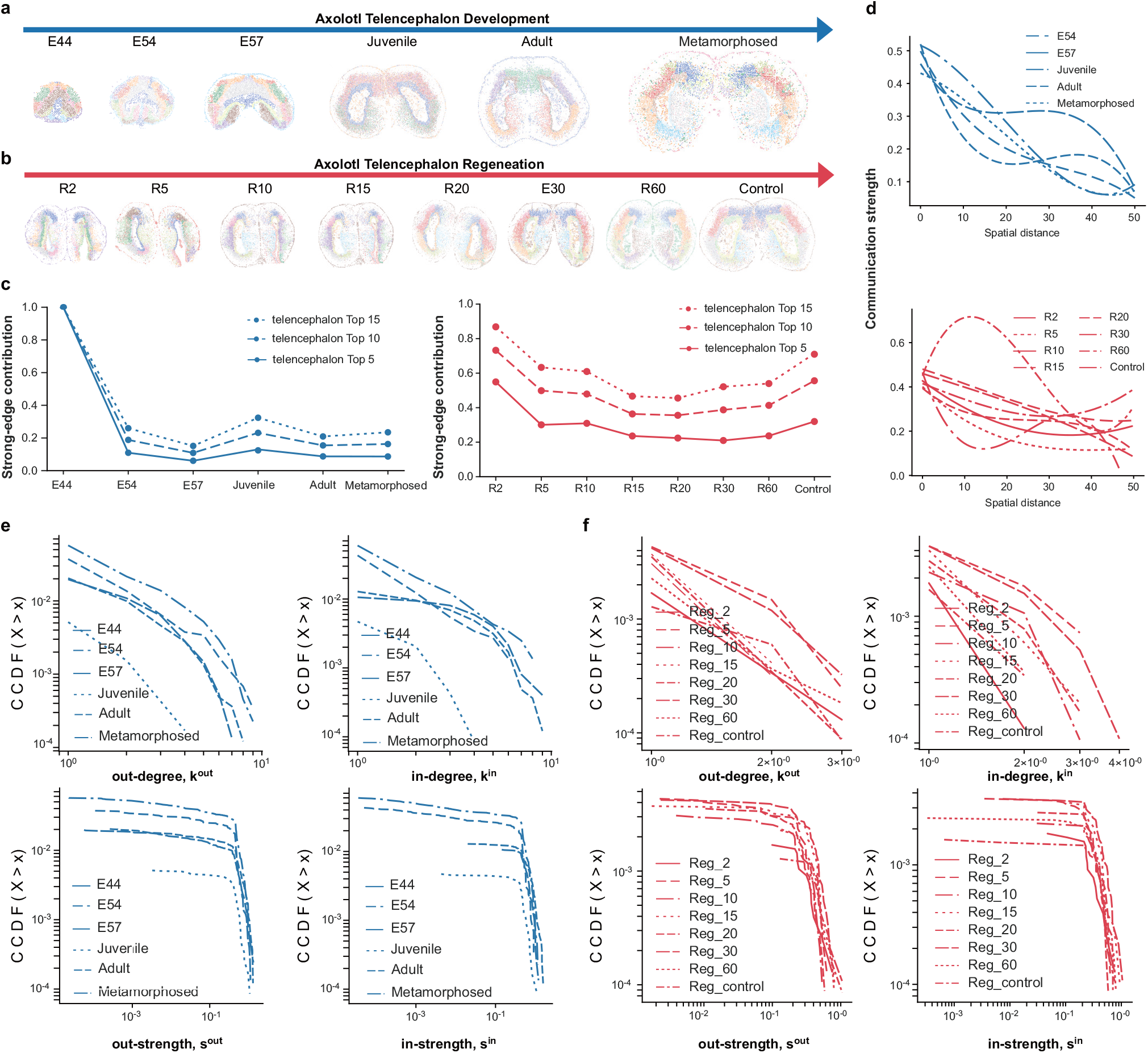
Additional topological characteristics of spatial cell–cell communication networks during axolotl telencephalon development and regeneration. **a**, Overview of the axolotl telencephalon developmental spatial transcriptomics dataset analysed in this study, showing representative sections across developmental stages. **b**, Overview of the axolotl telencephalon regeneration spatial transcriptomics dataset, showing representative sections across regenerative stages. **c**, Fractions of total communication contributed by the most highly connected nodes in the WNT7B-FZD5 communication network during axolotl telencephalon development (left) and regeneration (right), shown for the top 5, top 10, and top 15 nodes. **d**, Distance-dependent decay in communication strength for the WNT7B-FZD5 communication network during axolotl telencephalon development (top) and regeneration (bottom). **e–f**, Complementary cumulative distribution function (CCDF) curves of out-degree, in-degree, out-strength, and in-strength for the WNT7B-FZD5 communication network across axolotl telencephalon developmental **(e)** and regenerative **(f)** stages.

## Notes

### Competing Interest Statement

The authors have declared no competing interest.

### Summary of Updates

We have streamlined the Results section for clarity and conciseness by removing redundant text and improving the flow of the presentation. The analyses and conclusions remain unchanged.

